# Dendritic cell ICAM-1 strengthens immune synapses but is dispensable for effector and memory responses

**DOI:** 10.1101/2021.10.27.466043

**Authors:** Anita Sapoznikov, Stav Kozlovski, Sara W Feigelson, Natalia Davidzohn, Noa Wigoda, Ester Feldmesser, Ekaterina Petrovich-Kopitman, Anika Grafen, Moshe Biton, Ronen Alon

**Affiliations:** Dept. of Immunology, The Weizmann Institute of Science, Rehovot, Israel; Department of Biological Regulation, The Weizmann Institute of Science, Rehovot, Israel; Department of Life Sciences Core Facilities, The Weizmann Institute of Science, Rehovot, Israel; Department of Systems Immunology, The University of Würzburg, Würzburg, Germany

**Author notes:** Correspondence: Ronen Alon, Ph.D., Department of Immunology, The Weizmann Institute of Science, Rehovot, 76100, Israel.; Moshe Biton, Ph.D., Department of Biological Regulation, The Weizmann Institute of Science, Rehovot, 76100, Israel.

## Abstract

Lymphocyte priming in lymph nodes (LNs) depends on the formation of functional TCR specific immune synapses (ISs) with antigen (Ag) presenting dendritic cells. The high affinity LFA-1 ligand ICAM-1 has been implicated in different ISs studied *in vitro*. The in vivo roles of DC ICAM-1 in Ag stimulated T cell differentiation have been unclear. In newly generated DC conditional ICAM-1 knockout mice, we report that under Th1 polarizing conditions, ICAM-1 deficient DCs could not engage in stable conjugates with newly generated CD8 blasts. Nevertheless, these DCs triggered normal lymphocyte priming, proliferation and differentiation into functional cytotoxic T cells (CTLs) and central memory lymphocytes (Tcm) in both vaccinated and virus infected skin. Single cell RNAseq analysis confirmed that Tcm were normally generated in these mice and gave rise to normal T effectors during a recall skin response. Our results suggest that although CD8 T cell blasts tightly bind DC-ICAM-1, strongly adhesive DC-T ISs are not necessary for functional TCR dependent DC mediated CD8 T cell proliferation and differentiation into productive effector and memory lymphocytes.

**Summary:** Sapoznikov et al generated a new genetic murine model deficient in dendritic cell expression of the key adhesion molecule ICAM-1 and found that CD8 lymphocytes do not require strong adhesion to dendritic cells for antigen-dependent differentiation into effector and memory T cells.

## INTRODUCTION

Upon arrival to LNs, naïve T cells rapidly scan professional Ag-presenting cells (APCs), mainly classical DCs (cDCs) for cognate peptide-MHC complexes (*1, 2*). These DCs must provide naïve cells with an Ag-specific ‘Signal 1’, a co-stimulatory ‘Signal 2’, and a DC-secreted polarization ‘Signal 3’, in the form of cytokines (*3*). *In-vivo* multiphoton imaging suggests that soon after entry into LNs, CD8 T cells engage cDCs via multiple (serial) transient contacts (*1*) termed kinapses, before they generate long-lasting contacts (synapses) (*4, 5*). It is widely recognized that these stable ISs are necessary for full-fledged T cell differentiation (*6*), but the precise contribution of individual adhesion and costimulatory molecules expressed by different antigen presenting DCs to distinct types of ISs with different subsets of T cells during early exposure to antigenic moieties in LNs is still unclear. It was traditionally anticipated that long lived antigen-specific contacts between T cells and DCs requires integrin-mediated adhesions, in particular lymphocyte function-associated antigen 1 (LFA-1) mediated contacts (*5*), the involvement of this and other lymphocyte expressed integrins and their DC expressed ligands in T cell priming, proliferation, and differentiation into distinct subsets of effector and memory T cells is still unclear (*7*).

Intercellular adhesion molecule 1 (ICAM-1) is a ubiquitously expressed Ig superfamily high-affinity ligand of the lymphocyte integrin LFA-1 (*8–10*). The importance of LFA-1-ICAM-1 interactions for the formation of functional ISs has been demonstrated in many *in vitro* experimental settings, but rarely corroborated *in vivo* (*11–13*). Since genetic ablation of LFA-1 impairs T cell entry into LNs and Treg generation (*14, 15*) as well as homotypic T cell interactions (*16*), we previously addressed this question by following wild-type (WT) transgenic T cells in chimeric mice deficient in leukocyte ICAM-1 and ICAM-2 (*17*). We found that in a DC-targeted vaccination model in skin draining LNs triggered by intra-dermal injection of OVA combined with anti-CD40 mAb, DC ICAMs are dispensable for initial naïve CD4 T cell contacts with Ag-presenting DCs analyzed by intravital microscopy of skin draining LNs (*17*). In that study, the possibility that Ag activated CD4 T cells that differentiated into early T blasts use their LFA-1 to bind DC ICAM-1 and generate long-lasting ISs with Ag-presenting DCs was not investigated. That study also left open the possibility that ablation of DC expressed ICAMs might selectively affect subsequent differentiation processes taking place in the vaccinated LNs days after initial challenge (*6*). In addition, our previous study left unresolved the potential contribution of DC ICAMs to CD8 T cell activation and differentiation events taking place during viral infections (*18–21*). The maturation and reprograming of distinct skin migratory DCs and resident LN DCs vary between virus infected and differently vaccinated LNs (*22, 23*). Consequently, the differentiation of proliferating T cells into effector and memory subsets elicited by different vaccinations and infections involve distinct types of ISs.

The direct contribution of DC expressed ICAMs to T cell-DC contacts following vaccination and viral infection was traditionally dissected with adoptively transferred DCs (*12*). However, the properties and function of exogenous DCs (*24, 25*) do not fully recapitulate endogenous migratory and LN resident DCs. Furthermore, in total ICAM-1 knock out mice, the contribution of this key LFA-1 ligand expressed by DCs to distinct stages of T cell activation and differentiation taking place inside LNs cannot be distinguished from the contributions of ICAM-1 expressed by numerous other lymphocyte-educating LN stromal cells (*12*). These include vascular and lymphatic cells (*26, 27*), fibroblast reticular cells (FRCs) (*28*), follicular dendritic cells (FDCs) (*29–31*) and differentially activated T cells. ICAM-1 on subcapsular macrophages may also modulate the efficiency of viral antigen spread inside infected LNs (*32, 33*), and B cell expressed ICAM-1 (*34*) may contribute to CD4 T cell differentiation. We have therefore constructed a new strain of ICAM-1^fl/fl^ mice and generated DC ICAM-1 knockout mice by crossing the ICAM-1^fl/fl^ strain with mice expressing the Cre recombinase under the DC enriched CD11c promoter (*35–37*). We used these mice to follow the specific contributions of DC ICAM-1 to CD8 priming and differentiation into effector and memory CD8 T cells in distinct vaccinated and virus-infected LNs (*38, 39*). We chose naïve OT-I transgenic CD8 T cells to address these questions inside skin draining LNs following vaccination or skin infection with OVA-encoding modified vaccinia virus Ankara (MVA). We chose to address the role of DC ICAM-1 in CD8 rather than in CD4 T cell priming proliferation and differentiation in light of the much broader information gained so far on the former T cells (*1, 12, 18, 19, 40, 41*). To assess the role of DC ICAM-1 in stable ISs, we also made use of a recently developed flow cytometry based DC-T conjugate assay (*42*), which allowed us to quantify the formation of stable contacts between CD8 lymphoblasts and endogenous DCs recovered from vaccinated or virus infected skin draining LNs. We report that selective ablation of ICAM-1 from all endogenous cDCs and plasmacytoid DCs (pDCs) does not abrogate the ability of these DCs to mount normal OT-I CD8 lymphocyte priming and differentiation into effector and memory T subsets following skin vaccination and viral infection. Nevertheless, Ag presenting ICAM-1 deficient cDCs failed to generate stable conjugates with recently activated CD8 T cells. These unexpected results suggest that low stability ICAM-1-independent Ag-specific DC-T synapses are sufficient for full-fledged differentiation of naïve CD8 T cells into both effector and memory T cells.

## RESULTS

### ICAM-1 is selectively lost from all cDCs in both resting and vaccinated skin draining LNs of CD11C-Cre:ICAM-1^fl/fl^ mice

To dissect the function of DC ICAM-1 separately from the potential accessory roles of ICAM-1 expressed by other LN cells in T cell priming, we created a DC ICAM-1 knock out mouse by breeding CD11c-Cre mice with an in house generated ICAM-1^fl/fl^ mouse (Fig. S1 and Materials and Methods). We chose the CD11c Cre to silence ICAM-1 expression, since it has been extensively used for targeting various genes in different DCs (*35–37*). The major DC subset involved in CD8 priming, namely cDC1, lost the majority of their ICAM-1 in CD11c-Cre:ICAM-1^fl/fl^ mice in both resting as well as skin vaccinated popliteal (skin-draining) LNs (Fig. 1). cDC2, monocyte derived DCs (moDCs) and pDCs, accessory IFNγ producing DCs involved in Th1 and Tc1 differentiation processes also lost ICAM-1 expression (Fig. 1). The extent of ICAM-1 deletion in B cells and subcapsular sinus (SCS) macrophages, however, was negligible (Fig. 1). Notably, cDCs expressed residual, but comparable ICAM-2 expression in both control and CD11c-Cre:ICAM-1^fl/fl^ mice (Fig. S2). Thus, the main antigen presenting cDCs in our CD11c-Cre:ICAM-1^fl/fl^ mice (herein DC-specific ICAM-1 KO or DC^ΔICAM-1^ mice) are deficient in expression of the two main LFA-1 ligands ICAM-1 and ICAM-2 and could be compared side by side to cDCs from their Cre negative ICAM-1^fl/fl^ counterparts (herein referred to as control mice).

**Figure 1.**
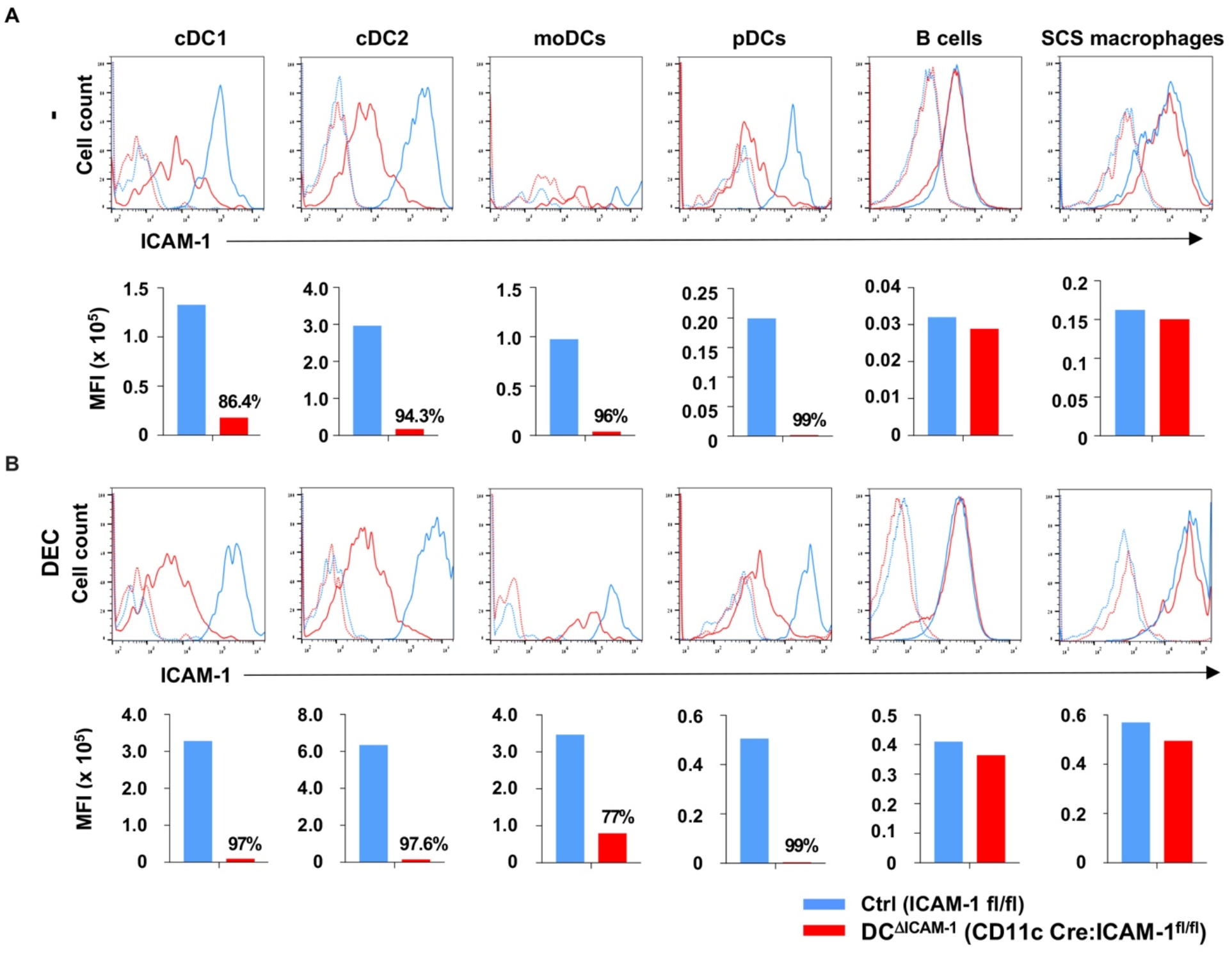
Selective deletion of ICAM-1 in DC subsets of popliteal LNs in conditional CD11C:ICAM-1 KO mice. Conditional CD11c:ICAM-1 KO (DC^ΔICAM-1^, red, solid line) or ICAM-1^fl/fl^ (control, blue, solid line) mice were injected intrafootpad with αDEC-205:OVA (0.05 µg/footpad) plus αCD40 (10 µg/footpad) (DEC) or PBS (-) and 24 hrs later different cell populations were recovered from popliteal LNs and analyzed for ICAM-1 expression. Isotype mAb controls, are shown in dashed lines. Bar graphs represent mean fluorescence intensity (MFI) of ICAM-1 expression in each of the indicated subsets. The extent of reduction of ICAM-1 expression on the indicated subsets is shown as % on top of the corresponding bars. The markers used to define each cell population are outlined in the Materials and Methods section. A representative of three experiments. At least 3 mice were analyzed for each group.

### DC ICAM-1 is dispensable for naïve CD8 activation and proliferation in DEC-205 targeted OVA-vaccinated skin draining LNs

The OVA antigen can be targeted into DCs residing in skin draining LNs, as well as to dermal DCs at the site of antigen administration via the DEC-205 scavenger receptor (*43*). DEC-205 receptor is similarly expressed by both control and ICAM-deficient DCs (*17*). Anti-DEC205 mAb chemically conjugated to OVA i.e., αDEC-205:OVA (*12, 43*) injected together with αCD40 activating mAb, a potent DC adjuvant, has been extensively used to follow the activation and differentiation of OVA-specific transgenic CD8 and CD4 T cells under Th1 and Tc1 polarizing conditions (*12, 17, 44*). To avoid overactivation of the naïve transgenic T cells, we injected intrafootpad a high sub-saturating amount of αDEC-205:OVA (Fig. S3), which induced robust and indistinguishable activation and proliferation of OT-I T cells adoptively transferred 18 hrs before the cocktail injection in both control and DC-specific ICAM-1 KO mice (Fig. 2A-D). Earliest OT-I T cell activation marked by acquisition of CD69 took place 6 hrs after αDEC-205:OVA and αCD40 injection (Fig. 2B). Subsequent OT-I activation marked by the CD25 surface marker was observed 24 hrs post αDEC-205:OVA and αCD40 injection (Fig. 2C), but at that time point, none of the activated OT-I T cells (T blasts) have divided, as was validated by CFSE dilution analysis (Fig. 2D). As expected, OT-I T cells transferred into mice deficient in DC ICAM-1 normally accumulated inside the skin draining popliteal LNs during this vaccination (Fig. S4A) but remarkably, underwent identical early (i.e. CD69 acquisition) and late (i.e., CD25 acquisition) activation (Fig. 2B,C, Fig. S4B). Furthermore, the kinetics of initial OT-I division (i.e., 30 hrs post vaccine injection), as well as subsequent proliferation, were essentially unaffected by ICAM-1 deficiency on all endogenous DC subsets (Fig. 2D). Notably, ICAM-1 deficient DCs expressed comparable levels of CD40, as well as of the main co-stimulatory molecules CD80 and CD86, as their control counterparts, ruling out compensatory overexpression of these DC activation molecules in ICAM-1 deficient DCs (Fig. S5). ICAM-1 deficient DCs were also confirmed to lack expression of Vascular Cell Adhesion Molecule 1 (VCAM-1) and did not exhibit elevated levels of prototypic cytokines involved in Tc1 differentiation (*17*). Thus, DC-expressed ICAM-1 seems dispensable for naïve OT-I activation and proliferation following skin vaccination with OVA and αCD40 mAb.

**Figure 2.**
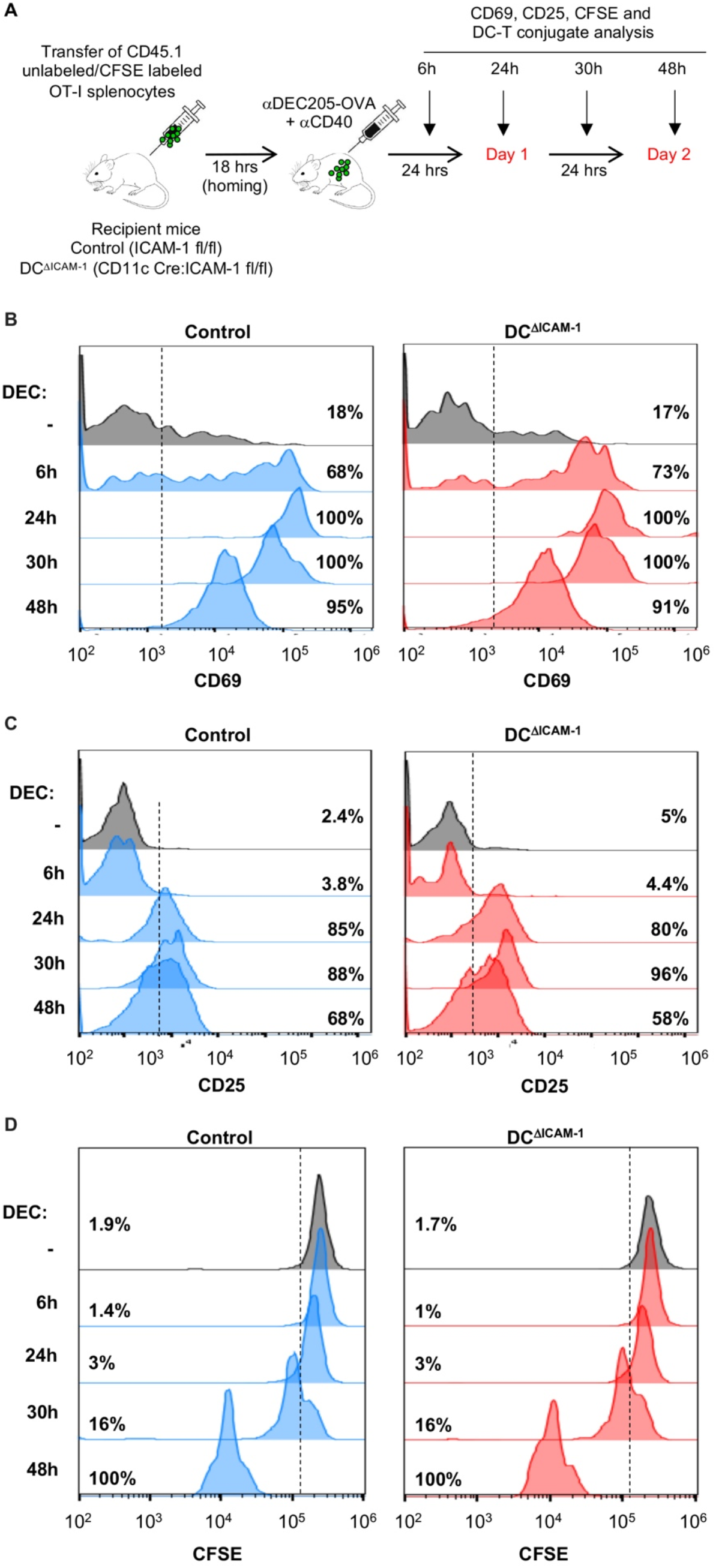
Normal early activation and first division of transferred OT-I T cells in αDEC205:OVA and αCD40-immunized LNs devoid of ICAM-1 expression on cDCs subsets. **(A)** A schematic representation of the experimental outline for Figure 2 and 3. CFSE-labeled OT-I CD45.1 T cells were adoptively transferred into control or DC^ΔICAM-1^ mice; 18 hrs later mice were immunized with intrafootpad injection of αDEC-205:OVA plus αCD40 and OT-I activation (CD69 and CD25), DC-OT-I cell conjugate formation and OT-I cell proliferation (CFSE), were assessed at the indicated time points. **(B)** Induction of CD69 or **(C)** CD25 on transferred naïve OT-I cells in popliteal LNs of either control (blue) or DC^ΔICAM-1^ (red) mice at indicated time-points after injection of αDEC205:OVA plus αCD40 (DEC) or PBS (-). **(D)** Proliferation histograms of CFSE-labeled OT-I T cells adoptively transferred into control (blue) or DC^ΔICAM-1^ (red) mice, and primed as indicated in A. CFSE levels and the percentage of CFSE-diluted lymphocytes were determined 6-48 hrs following immunization. At least three mice were analyzed in each experimental group.

### DC ICAM-1 is critical for rare stable Ag-dependent conjugates between cDCs and Ag-specific CD8 T blasts, but not for Ag stimulated DC-T clustering

Multiple multiphoton microscopy studies have indicated that Ag specific DC-T interactions in skin draining LNs follow a complex dynamics (*1, 6, 12, 40, 41*). Shortly after entering the T cell zone in these LNs, naïve T cells establish brief minute long contacts with multiple Ag presenting DCs (*1, 41*). Several hours later, a subset of activated T cells stop migrating and form stable long lived conjugates with single Ag presenting DCs (*6, 12*). We took advantage of a new flow cytometry-based conjugate assay (*42*), reported to detect stable contacts between naïve OT-1 cells and exogenous Ag presenting cDCs hours after intra-dermal vaccination with αDEC-205:OVA and αCD40 vaccine (*12*), and applied this technique to trace and quantify stable contacts between Ag activated OT-I T cells and endogenous cDCs in LN cell suspensions. In parallel, we used immunostaining to analyze DC-T clustering around DCs imaged in LN derived sections. In close agreement with the Scholer study, 24 hrs after intra footpad vaccination with αDEC-205:OVA and αCD40 cocktail, when all naïve OT-I T cells differentiated into OT-I T CD69+ CD25+ lymphoblasts, but have not yet divided, a fraction of these T blasts generated stable conjugates with endogenous LN cDCs (Fig. 3A, Fig. S6). These stable conjugates were enriched with LFA-1 on the T blasts engaging clustered ICAM-1 enriched on the DC surface (Fig. 3B). In agreement with the observations of Scholer *et al.* on exogenous mature DCs and adoptively transferred OT-I T cells (*12*), loss of ICAM-1 on all endogenous LN cDCs abrogated the formation of almost all stable DC and OT-I T cell conjugates recovered from the vaccinated LNs (Fig. 3C, left panel), whereas OT-I and DC numbers in both control and DC-specific ICAM-1 KO mice were similar (Fig. 3C, middle and right panels). Interestingly, immunofluorescence staining of endogenous cDCs and OT-I T blasts interacting inside vaccinated LNs indicated comparable numbers of blasts clustered around both control and ICAM-1 KO DCs (Fig. 3D,E). Importantly, at this time point, none of the blasts have divided (Fig. 2D). Thus, DC ICAM-1 appears critical for the ability of Ag stimulated OT-I T blasts to form rare firm conjugates with endogenous Ag presenting cDCs in vaccinated skin draining LN. However, DC ICAM-1 is dispensable for Ag stimulated T cell clustering around ICAM-1 deficient cDCs.

**Figure 3.**
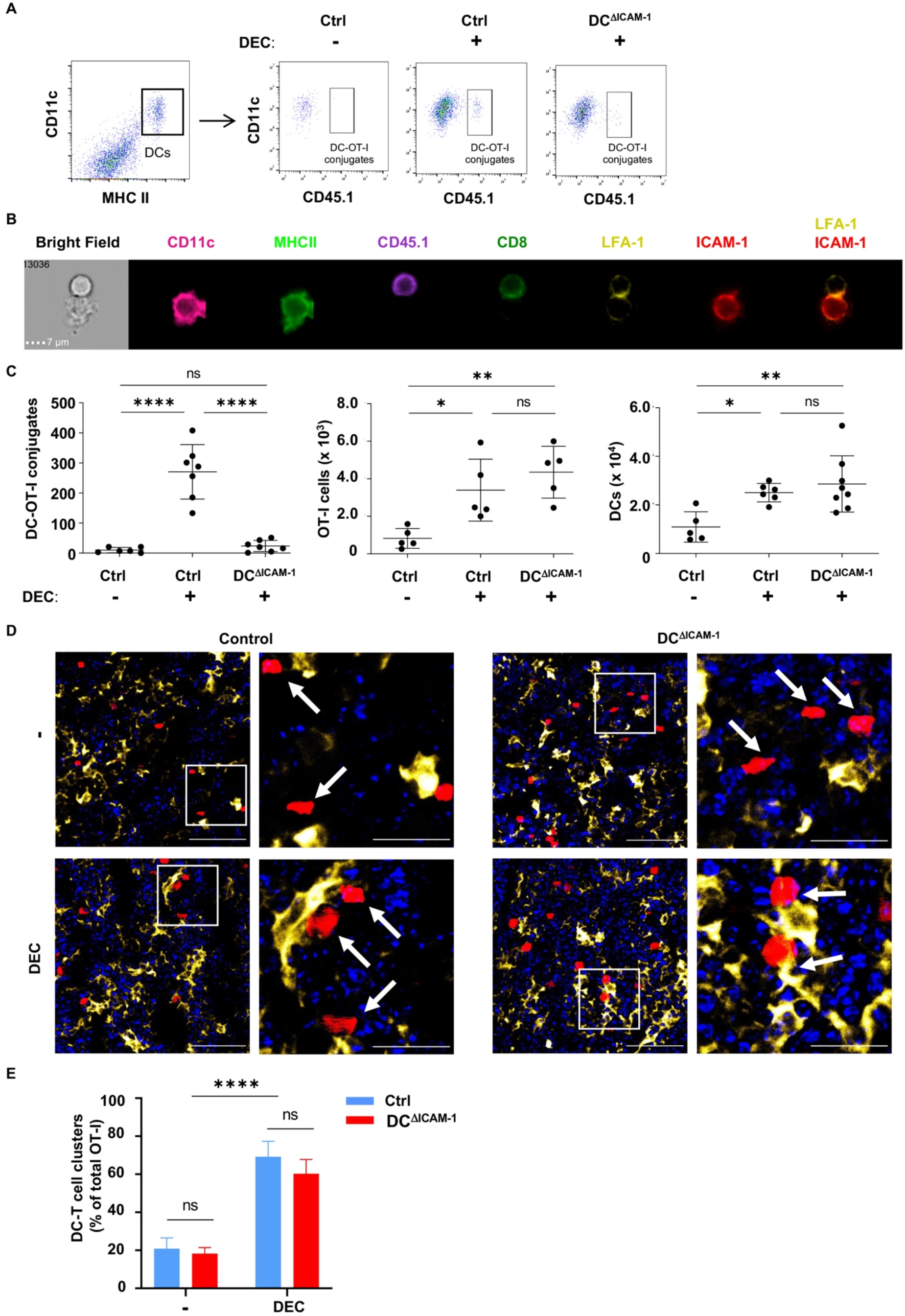
The loss of ICAM-1 on cDCs significantly reduces the frequency of stable DC-OT-I conjugates in immunized LNs without altering DC-T clustering. **(A)** Control or DC^ΔICAM-1^ mice were injected intrafootpad with αDEC205:OVA plus αCD40 18 hrs after OT-I CD45.1 intravenous transfer. DC-T cell conjugates were isolated 24 hrs later from popliteal LNs and analyzed by gating on doublets composed of cDCs (CD11c^high^ MHCII^high^ cells) and CD45.1 OT-I cells (CD3^+^, CD8^+^). Note, no conjugates in mice that were injected with PBS, and reduction in conjugate numbers in DC^ΔICAM-1^ versus control mice. **(B)** A representative image of a DC-T conjugate analyzed by image flow cytometry (ImageStream). The DC–T cell conjugate is depicted in the bright field and in individual fluorescent images. DCs were stained for CD11c and MHCII and OT-I cells were stained for CD3 (not shown), CD8 and CD45.1 markers. Note the accumulation of ICAM-1 and of LFA-1 at the contact area of the T cell and DC, respectively. *n* = 3. **(C)** Quantitative flow cytometric analysis of DC–T cell conjugates 24 hrs after vaccination with αDEC-205:OVA plus αCD40 (DEC) or PBS (-) in popliteal LNs of control and DC^ΔICAM-1^ mice (left). Total OT-I cells recovered from these LNs are shown in the middle panel and total DC numbers in the right panel. Each dot represents one animal. \**P* < 0.05, \*\**P* < 0.01 and \*\*\*\**P* < 0.0001 (one-way ANOVA with Tukey’s corrections). **(D)** Immunofluorescence staining of a popliteal LN section from control or DC^ΔICAM-1^ mice 24 hrs after intrafootpad injection either with αDEC205:OVA plus αCD40 (DEC) or PBS (-). 18 hrs prior to vaccination, tdTomato OT-I T cells (red) were intravenously transferred into the different recipient mice. DCs were stained with a directly labeled anti CD11c Ab (yellow). Cell nuclei were stained with Hoechst33342 (blue). Right panels depict magnified captions of the representative areas indicated by squares. Images are representative of six LNs isolated from three mice in each experimental group. Note the blastoid shape of the majority of T cells in the vaccinated LNs. Scale bars in the main figures are 50 µm and in the captions 20 µm. **(E)** Bar graphs represent the percentage of tdTomato OT-I T cells out of all OT-I T cells identified in each section which were in close contact with adjacent DCs upon vaccination as in (D). A total of 6 sections from 3 mice in each experimental group were analyzed. \*\*\*\**P* < 0.0001 (one-way ANOVA with Tukey’s corrections).

### DC ICAM-1 is not required for the generation of functional CTL effectors

We next followed the fate of the recently activated OT-I T cells in the LNs undergoing the standard αDEC-205:OVA and αCD40 vaccination in our conditional DC-specific ICAM-1 KO mice with respect to three critical parameters: T cell division and accumulation, differentiation into IFNγ, granzyme B and TNFα producing OT-I T cells, and differentiation into functional cytotoxic CD8 T cells that can *in vivo* kill Ag-loaded target cells inside the vaccinated LN. Remarkably, the numbers of Ag activated OT-I and their extent of proliferation that could be recovered at distinct time points following vaccination were not reduced in DC-specific ICAM-1 KO mice (Fig. 4A,B and Fig. S7). Thus, although facilitating stable Ag specific DC-T contacts, DC ICAM-1 was not essential for subsequent CD8 T proliferation induced by Ag and αCD40 stimulation. Surprisingly, however, and in contrast to the Scholer *et al.* study performed on similar LNs of total ICAM-1 KO mice (*12*), the number of effector OT-I T cells with high IFNγ, granzyme B and TNFα content in the absence of DC ICAM-1 was comparable to the controls (Fig. 4C, Fig. S8). Furthermore, DC ICAM-1 was dispensable for normal OT-I proliferation and differentiation triggered also by a 5-fold lower dose of αDEC-205:OVA (data not shown). Remarkably, the *in vivo* cytotoxic activity of these T cells evaluated by their capacity to eliminate SIINFEKL-loaded target cells reaching the vaccinated LNs 4 days post initial vaccination was also conserved in DC-specific ICAM-1 LNs (Fig. 4D,E). Thus, although DC ICAM-1 facilitates long lasting DC contacts with OT-I lymphoblasts 24 hrs post vaccination, these contacts and subsequent interactions of newly dividing OT-I T cells with DCs do not require the presence of DC ICAM-1 for the OT-I T cells to differentiate into fully functional cytotoxic T cells, as well as to effector T cells with high content of IFNγ and granzyme B. Since CD8 T_eff_ generated inside the vaccinated LNs egress these LNs and home to sites of infection, we next followed their ability to accumulate inside inflamed skin in a remote organ. To measure the migration capacity of T effectors generated in the vaccinated LNs of either control or DC-specific ICAM-1 KO mice, we followed the ability of OT-I T_eff_ endogenously stimulated by our vaccination protocol to enter a remote Complete Freud’s adjuvant (CFA) inflamed skin (e.g., ear dermis) 5 days after initial intrafootpad vaccination, independently of OVA presence at this skin site (Fig. 4F). As expected, naïve OT-I failed to accumulate at this site, further supporting the conclusion that all OT-I T cells accumulated inside the inflamed ear were T_eff_ cells generated inside the vaccinated skin draining LN. Strikingly, effector OT-I T_eff_ cells generated in vaccinated DC-specific ICAM-1 KO mice accumulated inside inflamed ear skin in numbers comparable to the OT-I T_eff_ generated in vaccinated control mice (Fig. 4G). Thus, the presence of DC-ICAM-1 is not essential for the ability of differentiated CD8 T cells stimulated by a skin vaccine to acquire full skin homing potential.

**Figure 4.**
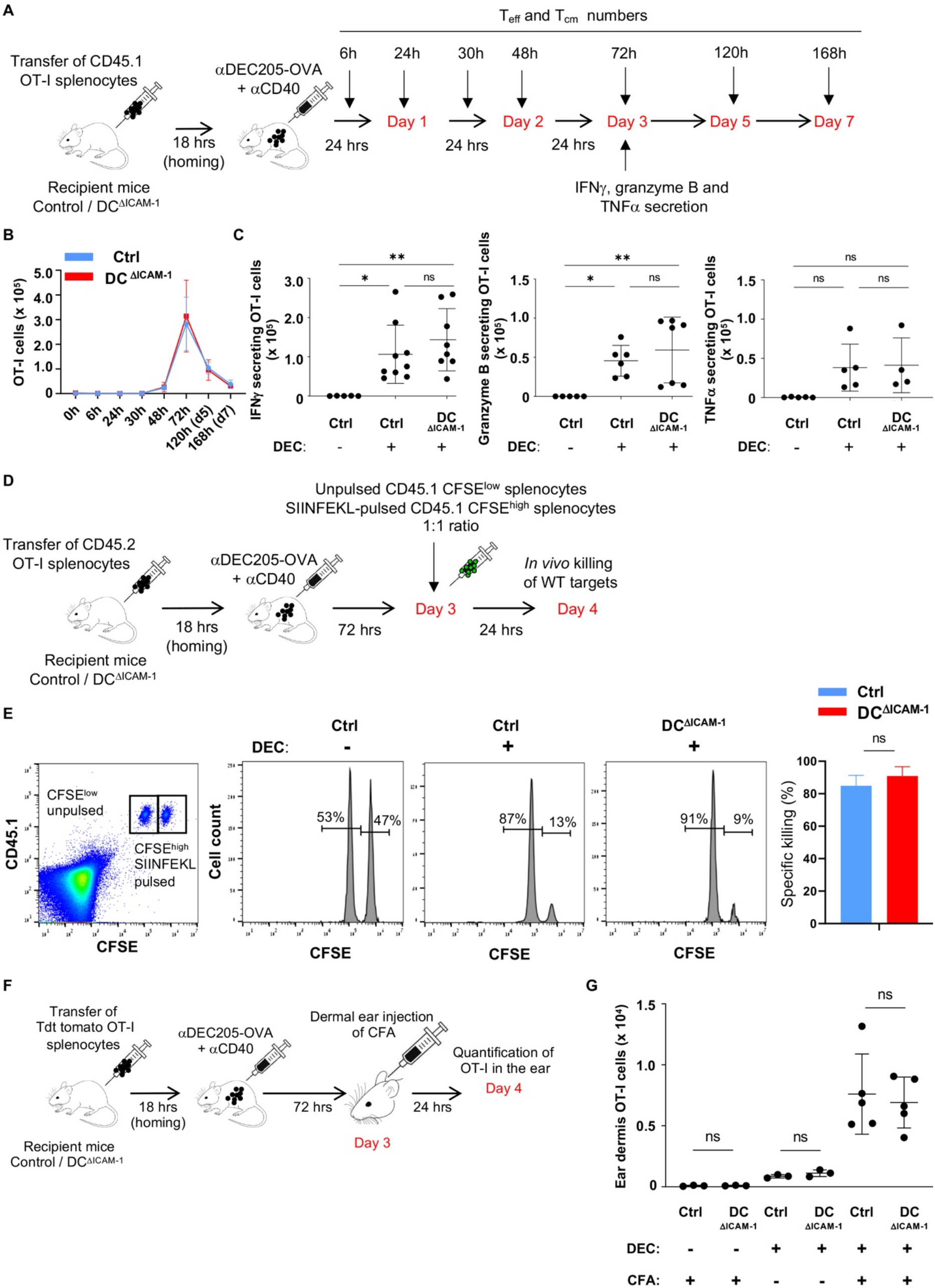
Normal differentiation and effector functions of OT-I lymphocytes generated inside vaccinated LNs of DC ICAM-1 depleted mice. **(A)** Control and DC^ΔICAM-1^ mice were injected intrafootpad with αDEC-205:OVA plus αCD40 18 hrs after OT-I CD45.1 transfer. The total numbers of OT-I cells recovered from popliteal LNs at 6-168 hrs following intrafootpad vaccination. IFNγ, granzyme B and TNFα staining of transferred OT-I T cells isolated from popliteal LNs was performed 72 hrs following vaccination. **(B)** The total numbers of OT-I cells recovered from popliteal LNs at the indicated time points following vaccination as shown in (A) in either control (blue, *n* = 3) or DC^ΔICAM^ (red, *n* = 7) mice. **(C)** Intracellular IFNγ, granzyme B and TNFα staining of transferred OT-I T cells isolated from popliteal LNs of control or DC^ΔICAM^ mice 72 h following vaccination. Each dot represents one animal. \**P* < 0.05 and (one-way ANOVA with Tukey’s corrections). **(D)** A scheme depicting the *in vivo* killing assay used to analyze *in situ* cytotoxic activity of differentiated OT-I T cells. Naïve OT-I cells were transferred into control or DC-specific ICAM-1 KO mice and 18 hrs later the mice were immunized with αDEC-205:OVA plus αCD40. Three days later, recipient mice were injected with equal numbers of CD45.1 CFSE^high^ OVA peptide-pulsed splenocytes and with unpulsed-CFSE^low^ cells. Antigen-specific killing was determined in the popliteal LNs of recipient mice 1 day later based on *in vivo* loss of CFSE^high^ target splenocytes. **(E)** The percentage of CFSE^high^ CD45.1 splenocytes pulsed with OVA peptide and unpulsed-CFSE^low^ cells allowed to measure CTL activity of OT-I T cells transferred and primed as explained in D. Bar graph indicates the percentage of target cell lysis inside control LNs (*n* = 7) or in LNs of DC^ΔICAM-1^ (*n* = 8) mice (unpaired *t* test). **(F)** The capacity of OT-I differentiated cells primed in skin draining LNs to egress and accumulate in a remote site of skin inflammation. Tdt tomato OT-I were transferred into control or DC^ΔICAM-1^ recipient mice immunized with αDEC-205:OVA plus αCD40 as in A. Three days later, recipient mice were subcutaneously rechallenged with 25 μg of complete Freud’s adjuvant (CFA) injected into each ear. **(G)** OT-I T cells were recovered from the ears of either control or DC^ΔICAM-1^ mice 1 day later. Each dot represents one mouse (one-way ANOVA with Tukey’s corrections).

### DC ICAM-1 is also not essential for OT-I differentiation into functional central memory CD8 lymphocytes

Proliferating T cells differentiate into two main CD44 high subsets, distinguished by their levels of CD62L (L-selectin), namely, short lived effector T cells (CD62L^lo^) and long lived central memory (CD62L^hi^) subsets (*45, 46*) (Fig. 5A) that recirculate among LNs searching for signs of infection with pathogens expressing their cognate antigen. We therefore next compared the differentiation of adoptively transferred naïve OT-I into either T_eff_ or T_cm_ subsets at different time points following the intrafootpad vaccination with the αDEC-205:OVA and αCD40. Surprisingly, both T_eff_ and T_cm_ subsets were generated from naïve OT-I T cells at similar numbers in popliteal LNs of control and DC-specific ICAM-1 KO mice at different time points following vaccination (Fig. 5B). These results suggested that the presence of DC ICAM-1 is not obligatory for the proliferation and differentiation of naïve CD8 into both effector and central memory subsets.

**Figure 5.**
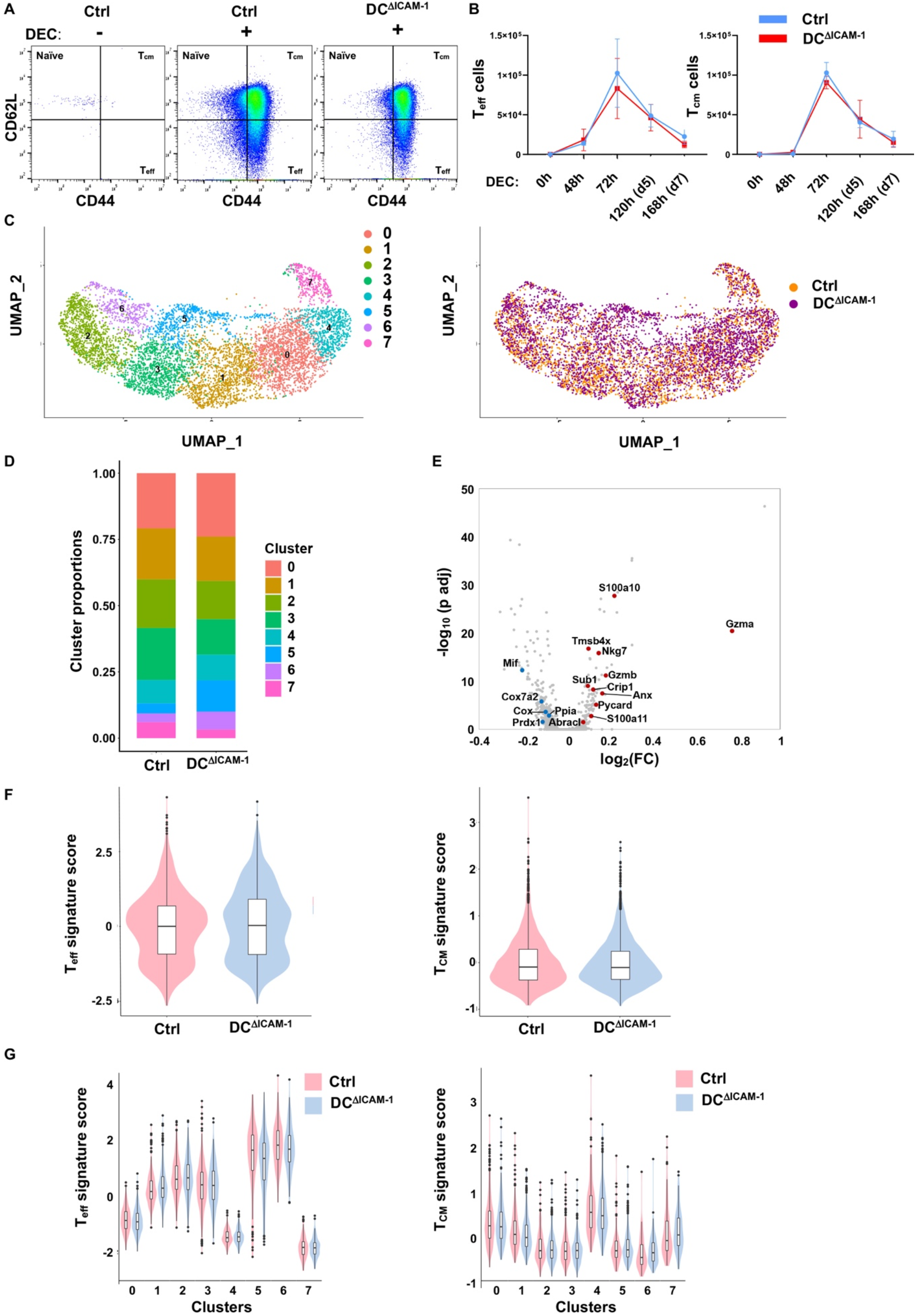
DC ICAM-1 is not obligatory for OT-I differentiation into effector and central memory CD8 lymphocytes. **(A)** Flow cytometry of CD3^+^ CD8^+^ OT-I T cells in popliteal LNs of PBS injected or 72 hrs after αDEC-205:OVA plus αCD40 intrafootpad immunization of control and DC^ΔICAM-1^ mice. T_cm_ (CD62L^+^ CD44^+^), T_eff_ cells (CD62L^-^ CD44^+^) and naïve cells (CD62L^+^ CD44^−^) can be readily distinguished. **(B)** The number of T_eff_ (left panel) and T_cm_ cells (right panel) recovered from control (blue, *n* = 3) or DC^ΔICAM-1^ (red, *n* = 7) mice at the indicated time-points following αDEC-205:OVA plus αCD40 immunization (DEC). (**C-G**) Single-cell analysis of CD8 OT-I cells isolated from αDEC-205:OVA plus αCD40 immunized control or DC-specific ICAM1 deficient mice. (**C**) OT-I cell subsets. Uniform Manifold Approximation and Projection (UMAP) for Dimension Reduction of 6993 single cells (points), colored by cluster assignment (left panel) or by host genotype (right panel), n=2 samples per genotype. (**D**) Proportion of different OT-I subsets (colored), as determined by scRNA-seq of 6993 cells sorted from control or DC-specific ICAM-1 KO mice (n=2 per group). (**E**) Volcano plot of DE genes based on 3814 OT-I cells isolated from DC ICAM-1 KO mice LNs (n=2 mice) vs. 3179 OT-I cells from matched control LNs (n=2 mice). Red dots: upregulated T_eff_ genes, blue dots: downregulated T_eff_ genes (FDR < 0.05, likelihood-ratio test), gray dots: all other genes. (**F,G)** Expression of effector and memory T cell signatures in OT-I T cells. Violin plots showing the distribution of mean expression values of T_eff_ signature genes (left, top 20 genes,(*81*)) or T_cm_ signature genes (right, top 20 genes, (*81*), see Materials and Methods) isolated from immunized control or DC-specific ICAM-1 KO mice (*n*=2 mice per group). **(F)**, all OT-I cells, **(G)**, OT-I T cell subsets. No significant statistical changes were found (FDR). Bar showing median and box interquartile range.

To further address the possibility that DC-ICAM-1 contributes to specific Ag driven differentiation programs of T effectors and central memory T cells, we next dissected the transcriptional programs of sorted adoptively transferred OT-I T cells from control or DC-specific KO mice single cell RNA sequencing (scRNA-seq). The OT-I T cells were isolated from the αDEC-205:OVA and αCD40 vaccinated skin draining LNs 3 days after initial priming, a time point of peak accumulation of both effector and memory Ag-specific T cells in these LNs (Fig. 5B). Sorted OT-I cells from both control and DC-specific KO mice were labeled with hashed-tagged antibodies to minimize batch effects (Fig. 5C and Fig. S9A-D). Single cell transcriptomic analysis revealed 8 OT-I T cell subsets with similar proportions (see Table S1), but with a slight increase in predicted T_eff_ frequency (clusters 5 and 6, Fig. 5D,G) of the DC-specific ICAM-1 KO mice. These data suggest that CD8 T cell interacting with Ag presenting ICAM deficient DCs readily differentiate into normal effector T cells (Fig. 5D,F,G). Furthermore, the single cell RNA analysis indicated strikingly similar transcription patterns of predicted memory OT-I T cells in vaccinated control versus DC-specific ICAM-1 KO LNs (cluster 4, Fig. 5D,F,G). Genes upregulated during transition and maintenance of memory CD8^+^ T cells (*47*) like *Il7r*, *Sell* (L-selectin) as well as transcriptions factors (*Runx1*, *Prdm1*) and co-stimulatory molecules (*Ctla4*, *Icos*) were also largely conserved between the compared experimental groups (see Table S2). Surprisingly, a small subset of effector OT-I generated in DC-specific ICAM-1 KO LNs expressed elevated levels of molecules associated with highly activated CTLs (e.g., *Gzma*, *Gzmb* and *Ccl5*, Fig. 5E and Fig. S9E,F). As expected for this vaccination, exhaustion markers such as *Havcr2* (Tim3) and *Pdcd1* (PD-1) were either absent or low in both OT-I groups (see Table S2). Furthermore, the profiles of canonical inhibitory signature reported for effector CD8 T cells (*48*) were also conserved for OT-I isolated from the vaccinated control or DC-specific ICAM-1 KO LNs (Fig. S9G,H). Collectively, the single cell analysis corroborated with our functional assays and indicate that ICAM-1 is not essential for robust vaccination driven DC-mediated CD8 T cell differentiation into effector and memory T cells in vaccinated LNs.

In order to assess whether the central memory T cells generated during the vaccination protocol in our DC-specific ICAM-1 KO mice are fully functional as suggested by the single cell transcriptomic analysis, we next exposed the OT-I T cells primed by the αDEC-205-OVA and αCD40 vaccine to a secondary skin challenge 12 days later, introduced by an intrafootpad injection of OVA in the presence of the potent adjuvant CFA (Fig. 6A). Strikingly, 4 days after the recall OVA/CFA immunization, the numbers of OT-I T cells recovered from the immunized skin draining LNs and the relative fraction of IFNγ, granzyme B and TNFα high T effectors were comparable in control and DC-specific ICAM-1 KO mice (Fig. 6B). Since without a recall challenge, no OT-I effectors could be recovered from LNs at this time point (Fig. 6A), suggesting that the pool of originally generated T_eff_ and T_cm_ cells had contracted or egressed from the vaccinated LNs (Fig. 4B, 5B). These effectors were most likely generated from T_cm_ subsets exposed to the recall challenge in the draining LNs as previously reported (*49–51*). Collectively, these results suggest that the OT-I T_cm_ generated during the primary response in the absence of DC ICAM-1 (Fig. 5A,B,G) homed normally to inflamed skin draining LNs during a recall skin response, where they underwent effective recall activation and differentiation in the absence of DC ICAM-1 in the rechallenged skin draining LNs. To further prove that the T_cm_ generated in vaccinated skin draining LNs of DC-specific ICAM-1 KO mice are as functional as those generated in vaccinated skin draining LNs of control mice, we next isolated sorted T_cm_ from the vaccinated LNs of either control or DC-specific ICAM-1 KO mice (recipient I, Fig. 6C) and transferred these T_cm_ into WT recipient mice (recipient II, Fig. 6C). In this configuration, the T_cm_ entering the skin draining LNs of OVA/CFA challenged mice underwent *in situ* activation and differentiation in the presence of control ICAM-1 expressing DCs. Consistent with the results described in Fig. 6B, the adoptively transferred T_cm_ derived from either control or DC-specific ICAM-1 KO mice gave rise to comparable numbers of IFNγ, TNFα and granzyme B rich T effectors in the OVA/CFA challenged LNs of WT recipient mice (Fig. 6D). Since long lasting OT-I memory was severely perturbed in naïve OT-I generated in similarly vaccinated skin draining LNs of total ICAM-1 KO mice (*12*), our results suggest that OT-I memory responses depend on ICAM-1 expression by accessory LN cells other than DCs.

**Figure 6.**
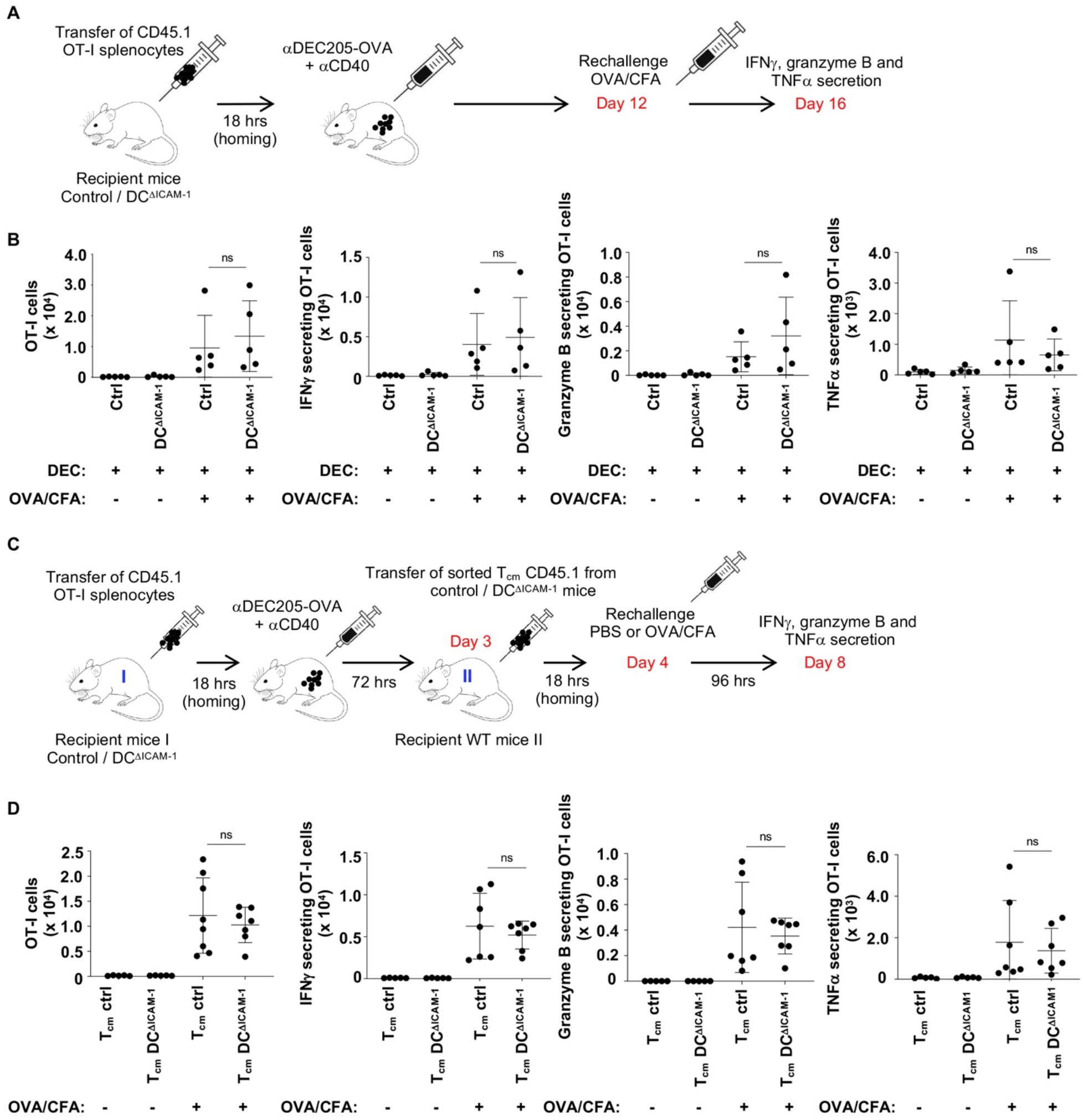
DC ICAM-1 is not obligatory for rapid generation of T effectors during a recall response following vaccination. **(A)** Mice were first intrafootpad immunized with αDEC-205:OVA plus αCD40 (18 hrs post intravenous OT-I transfer), and 12 days later were subcutaneously boosted with 40 µg of OVA protein in CFA or PBS. LNs were harvested and analyzed 4 days after the boost. **(B)** Quantification of effector OT-I T cell numbers and their intracellular IFNγ, granzyme B and TNFα staining in popliteal LNs of either control or DC^ΔICAM-1^ mice subjected to a recall immunization. Each dot represents a mouse (unpaired *t* test). **(C)** A scheme depicting recall response of OT-I T_cm_ sorted from control or DC^ΔICAM-1^ mice (I, blue) 3 days after immunization with αDEC-205:OVA plus αCD40. 18 hrs after T_cm_ transfer, WT mice (II, blue) were rechallenged subcutaneously with 40 µg of OVA protein in CFA or PBS and 4 days later, OT-I cell numbers and their intracellular IFNγ, granzyme B and TNFα staining were quantified in popliteal LNs **(D)**. Each dot represents one mouse (unpaired *t* test).

### DC ICAM-1 stabilizes MVA-triggered DC conjugates with Ag-specific T blasts, but is not required for subsequent proliferation and differentiation into functional effector and memory T cells

To further explore the contribution of DC ICAM-1 to OT-I differentiation following viral infection, we next challenged the OT-I T cells in our mice models post infection with a modified replication-deficient OVA encoding MVA, a widely used model of acute viral infection of skin draining LNs (*47, 52, 53*). We first validated that ICAM-1 expression in all major DC subsets involved in CD8 priming and differentiation during this viral infection, including pDCs, was significantly reduced following intrafootpad injection with MVA-OVA of the DC-specific ICAM-1 KO mice (Fig. 7A). Reminiscent of the vaccination results, OT-I T cells accumulated in the popliteal LNs of either control or DC-specific ICAM-1 KO recipient mice underwent identical OVA specific activation (Fig. S10). Interestingly, 24 and 30 hrs after intrafootpad viral injection, all Ag activated OT-I T lymphoblasts that acquired high levels of CD69 and CD25 (Fig. S10) have not divided yet (Fig. 7B). Although 30 hrs post infection, OT-I T blasts failed to generated stable conjugates with LN DCs recoverable from total LN suspensions of DC-specific ICAM-1 KO mice, at this time point a fraction of similar Ag-activated OT-I blasts stably engaged with control DCs (Fig. 7C). As observed in LNs vaccinated with the αDEC-205-OVA and αCD40 vaccine (Fig. 3B), these conjugates were enriched with a typical cluster of LFA-1 on the T blasts engaging clustered ICAM-1 on control DCs (Fig. 7D). Reminiscent of the conjugates generated following αDEC-205-OVA and αCD40 vaccination and while the numbers of OT-I T cells and DCs were undistinguishable in both control and DC-specific ICAM-1 KO mice at 30 hrs post infection (Fig. 7E, middle and right panels), the frequency of stable conjugates generated by OT-I blasts with endogenous ICAM-1 deficient DCs was substantially lower than with control DCs (Fig. 7E, left panel).

**Figure 7.**
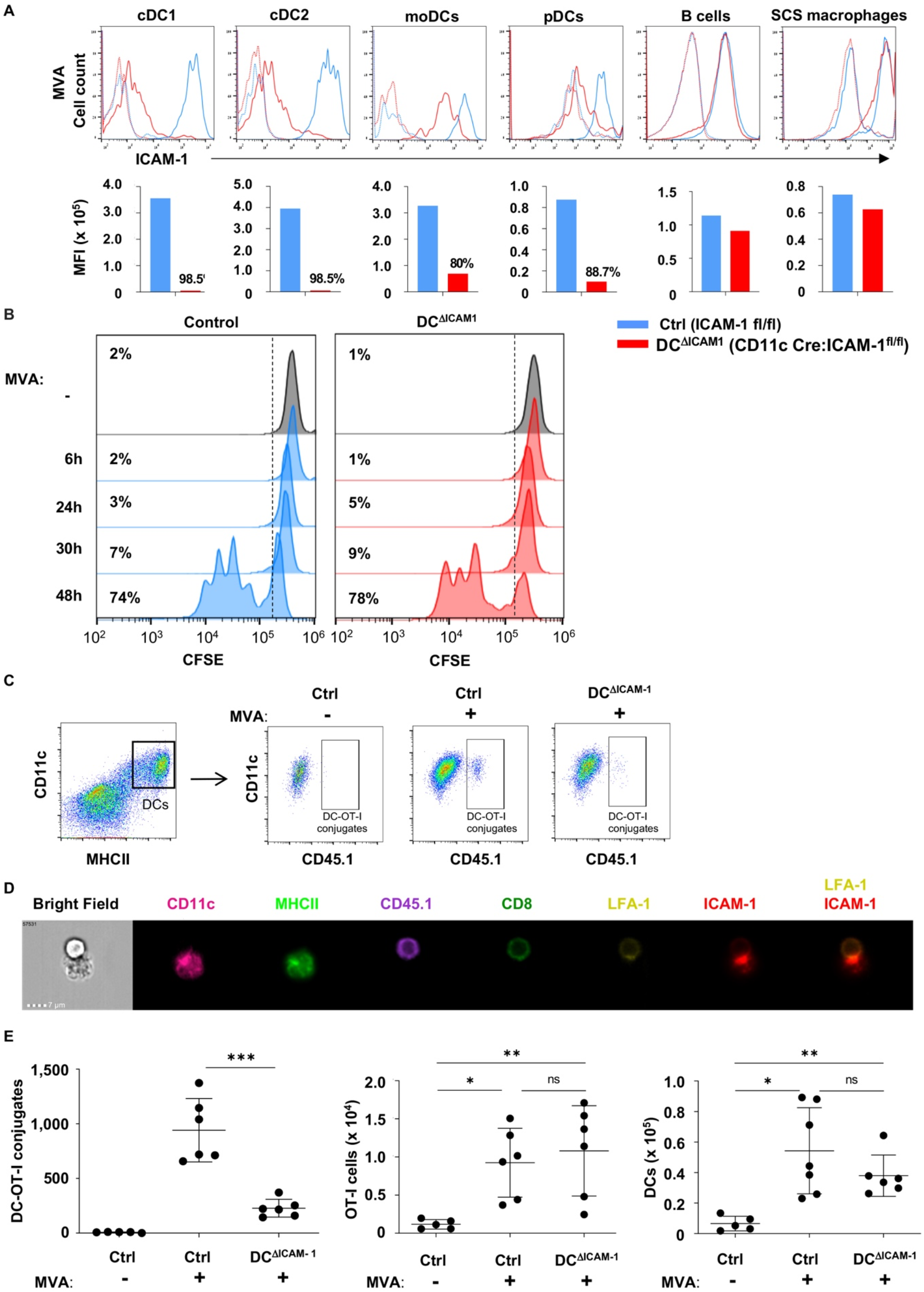
DC ICAM-1 is critical for stable formation of conjugates with Ag-specific CD8 T blasts triggered by MVA-OVA infection, but is not crucial for CD8 T cell proliferation, differentiation and recall response in the LNs. **(A)** Control and DC^ΔICAM-1^ mice were injected intrafootpad with MVA-OVA (1×10^7^ infection units, IU pe footpad, MVA) and 24 hrs later different cell populations were recovered from popliteal LNs and analyzed for ICAM-1 expression as in Figure 1. At least 3 mice were analyzed for each group. **(B)** Proliferation histograms of CFSE-labeled OT-I CD8 T cells adoptively transferred into control or DC^ΔICAM-1^ mice, primed 18 hrs later by intrafootpad injection of MVA-OVA (1×10^7^ IU/footpad) or PBS. Percentage CFSE-diluted cells was determined 6, 24, 30 and 48 hrs following infection. At least 3 mice were analyzed for each group. **(C)** Control and DC^ΔICAM-1^ mice were infected intrafootpad with MVA-OVA (1×10^7^ IU) 18 hrs after OT-I CD45.1 transfer. DC-T cell conjugates were isolated from popliteal LNs 30 hrs later and analyzed as explained in Fig. 3. **(D)** A representative DC-T cell conjugate analyzed by image flow cytometry. Note accumulation of ICAM-1 and LFA-1 at the contact area of the DCs and T cells. Data were collected from 3 mice. **(E)** Quantitative flow cytometric analysis of the number of recovered DC–T cell conjugates (left), OT-I T cells (middle) and DCs (right) in popliteal LNs of control and DC^ΔICAM-1^ mice 30h following MVA-OVA infection (1×10^7^ IU). Each dot represents one mouse. \**P* < 0.05, \*\**P* < 0.01 and \*\*\**P* < 0.001 (one-way ANOVA with Tukey’s corrections).

We next tested whether effector and memory OT-I T cells are generated at different efficiencies in the absence of cDC and pDC expressed ICAM-1. Strikingly, the numbers of Ag activated OT-I that underwent subsequent proliferation and differentiation into effector IFNγ rich CD8 T cells following MVA-OVA injection were not reduced in the absence of cDC and pDC ICAM-1 (Fig. 8A and Fig. S11). Furthermore, effective memory responses were observed during a recall OVA/CFA immunization 16 days post viral infection: indeed, 4 days after a footpad immunization with OVA/CFA, the total numbers of OT-I cells and the relative fraction of IFNγ, granzyme B and TNFα high OT-I T effectors accumulated in rechallenged skin draining LNs were comparable in both infected control and DC ICAM-1 KO recipient mice (Fig. 8B). Interestingly, even without a recall response and in contrast to the vaccination results (Fig. 6B), residual effector OT-I T cells could be recovered from skin draining LNs 16 days post MVA-OVA infection (Fig. 8B, two first bars in each scheme). These residual OT-I effectors also did not require DC ICAM-1 to prevail in the MVA infected LNs during the contraction phase. Collectively our results suggest that the presence of ICAM-1 on LN cDCs and on pDCs during a primary viral infection, as well as during a recall response is dispensable for the generation of T_eff_ and T_cm_ cells capable of differentiating into newly generated T effectors in rechallenged skin draining LNs.

**Figure 8.**
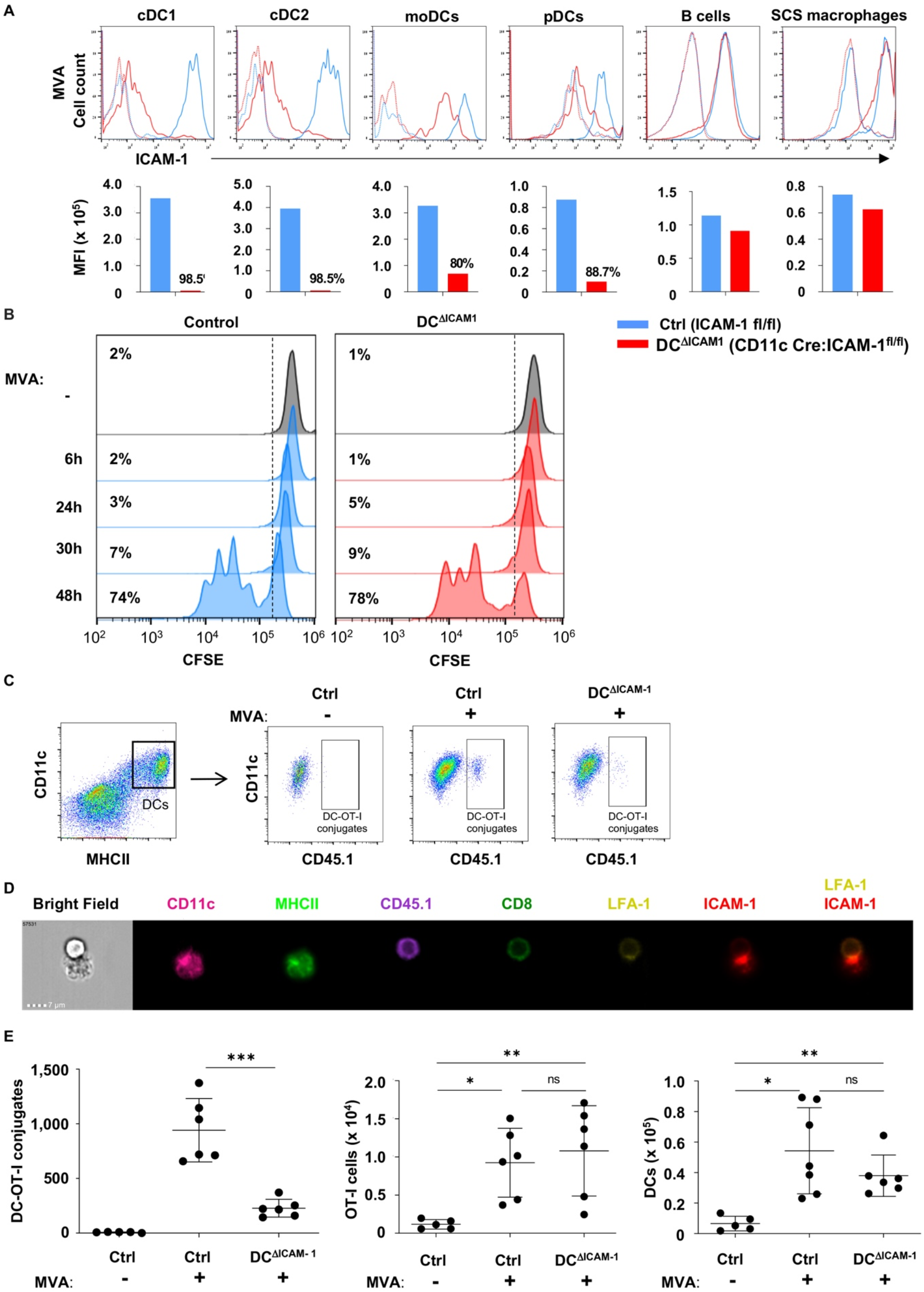
ICAM-1 expression on cDCs and pDCs is not crucial for CD8 T cell proliferation, differentiation and recall response in MVA-OVA infected LNs. **(A)** The total numbers of OT-I cells and of IFNγ producing OT-I cells recovered from LNs 72h following intrafootpad infection with MVA-OVA (1×10^7^ IU) of control and DC^ΔICAM-1^ mice. Each dot represents one mouse. \**P* < 0.05 and \*\**P* < 0.01 (one-way ANOVA with Tukey’s corrections). **(B)** Quantification of memory OT-I T cells and their intracellular IFNγ, granzyme B and TNFα stainings in popliteal LNs of control and DC^ΔICAM-1^ recipient mice. The mice were first intrafootpad injected with MVA-OVA (1×10^7^ IU) and 12 days later were subcutaneously boosted with PBS or 40 µg of OVA in CFA. LNs were analyzed 4 days after the rechallenge as outlined in Figure 6A. Each dot represents one animal (unpaired *t* test).

## DISCUSSION

The putative importance of LFA-1-ICAM-1 interactions for T cell binding of Ag presenting DCs and in the formation of functional immunological synapses with these professional APCs has been demonstrated in several *in vitro* experimental settings (*54, 55*), but rarely dissected *in vivo* (*12, 13, 17, 56–58*). Although the critical role of LFA-1, the main counter-receptor of DC ICAM-1 in T cell homing, proliferation or differentiation, was demonstrated in several *in vivo* studies (*54–56*), a direct role for DC ICAM-1 could not be derived in these and other studies due to promiscuous contributions of the LFA-1 ligands ICAM-1 and ICAM-2 to T cell engagements of FRCs, activated B cells as well as of activated T cell subsets (*59*). A previous comprehensive study highlighted the importance of LFA-1-ICAM-1 interactions of T cells with DCs for naïve CD8 differentiation and memory acquisition, but could not assign these processes directly to ICAM-1 expression specifically on DCs due to lack of an appropriate genetic model (*12*). To identify the direct role of DC ICAM-1 in initial T cell priming and differentiation, and distinguish its functions in DC mediated education of T cells from other co-stimulatory roles of ICAM-1 expressed by other LN cells, we have generated new DC ICAM-1 knockout mice, in which ICAM-1 was depleted primarily in all cDCs, as well as on pDCs via a CD11C driven Cre recombinase. In the present study, we used this new genetic system to follow if and how OT-I CD8 priming in skin-draining LNs under Th1 and Tc1 polarizing conditions implicates ICAM-1 expressed by lymph node DCs (*60*). Using a flow cytometry based assay for isolation of stable *in vivo* Ag driven DC-T conjugates from immunized LNs (*42, 61*), we have identified an early time window in which these conjugates could be isolated from either vaccinated or virus infected LN cell suspensions. Surprisingly, although these rare stable DC-T conjugates depended on DC expression of ICAM-1, endogenous Ag stimulated T cell clustering around DCs, depicted by histological analysis of intact LN sections, did not. These stable ICAM-1 strengthened conjugates between T blasts and Ag presenting DCs appear dispensable for subsequent T cell proliferation and differentiation inside vaccinated or infected LNs. These findings thus indicate for the first time that the strength of TCR-dependent T cell adhesion to Ag presenting DCs does not correlate with the efficiency of subsequent T cell activation by these DCs.

Previous studies have suggested that prolonged synapses between T blasts and Ag-presenting DCs vary with the strength of the integrated TCR signals, and depend on both the affinity of the pMHC and its relative density on the DC surface (*6*). Whether DC ICAM-1 can prolong TCR triggered T cell contacts by strong adhesive interactions, which could thereby facilitate continuous TCR signaling within DC-T synapses has been a long-standing question. Our new results are the first to suggest that while DC ICAM-1 can indeed promote tight T cell adhesions to DCs during Ag stimulated activation of naïve CD8 lymphocytes inside immunized LNs, the presence of this major LFA-1 ligand is in fact dispensable for the ability of Ag stimulated T blasts to cluster with DCs in response to proliferative antigenic signals and for the ability of additional Ag presenting cDCs and pDCs to provide subsequent differentiation signals to these blasts and to their daughter cells. Thus, ICAM-1 independent clustering of newly activated T cells with Ag presenting DCs, potentially via CD40-CD40L interactions (*62*), rather than tight ICAM-1 mediated adhesions, appears to be both required and sufficient for T cell proliferation and differentiation into both T_eff_ cells and T_cm_ in both vaccinated and virus infected skin draining LNs. Notably, the T_eff_ generated in our DC-specific ICAM-1 KO mice were not only highly cytotoxic, but once egressed from immunized LNs, accumulated normally at effector sites in the skin. These results further suggest that as part of their differentiation inside skin draining LNs, the imprinting of CD8 effectors with trafficking molecules critical for their homing to sites of skin infection does not require T cell engagement with DC ICAM-1. Furthermore, the T_cm_ generated in the DC specific ICAM-1 KO mice underwent normal *in vivo* expansion into T_eff_ cells in draining LNs during a recall response, further suggesting that DC-ICAM-1 is also dispensable for the Ag-driven transition of memory T cells into T_eff_ cells in these lymphoid organs. Nevertheless, a key open question that remains open is the long-term fate of these and other T_cm_ that are generated during distinct primary immunizations in the absence of DC ICAM-1, especially during viral infections. Notably, endogenous CD8 T cells deficient in LFA-1 exhibit reduced differentiation into effector T cells following infection with Listeria monocytogenes (*63*), but can still differentiate into functional T_cm_ indicating that T_cm_ can mount a robust secondary response to infection in the absence of LFA-1 (*63*). Our present results in mice lacking DC ICAM-1 are consistent with these results because upon a rechallenge, Ag specific T_cm_ generated in our DC-specific ICAM-1 KO mice and adoptively transferred into mice with normal ICAM-1 levels underwent normal differentiation into newly expanded T_eff_. Future longer term cell fate studies using similar adoptive transfer approaches of similar T_cm_ into WT recipients followed by their re-challenge will be required to further substantiate our present findings.

In a previous study using chimeric mice, we reported that CD4 OT-II T cells do not use DC ICAMs for initial priming, for Ag-TCR triggered semi-stable arrests (kynapses), and for generation of short lived T effectors (*17*). In that study the contribution of DC ICAMs for stable DC-T conjugates generated by T blasts was not investigated. Here, using a similar vaccination approach we found that DC ICAM-1 contributes to highly stable, but rare conjugates generated between CD8 T cells and Ag presenting DCs *in situ* stimulated with anti-CD40. The earliest DC ICAM-1 dependent DC-T conjugates are generated just prior to the initial T cell division into two daughter T cells. Why then are these stable DC-T conjugates dispensable for T cell proliferation and differentiation? Unexpectedly, high-resolution single cell transcriptomic analysis of these T effectors generated in the absence of DC ICAM-1 revealed a small subset of hyperactivated T effectors. Importantly, these and all other effector T cells differentiated in the absence of DC ICAM-1 effectively killed target cells *in situ*, egressed the LNs and homed into secondary sites of skin infection similarly to T effectors generated in WT LNs. We thus favor the possibility that in the setup of intradermal injected αDEC-205:OVA and αCD40, even at low doses of soluble DEC-205 targeted antigen, numerous OVA peptide/MHC presenting DCs, each stimulated with the anti CD40 mAb, can promote frequent DC contacts with either Ag stimulated CD4 or CD8 T cells that compensate for the reduction in the strength of individual CD4 or CD8 T interactions with small subsets of ICAM-1 deficient DCs. Thus, frequent low strength synapses between Ag presenting DCs generated during both vaccination and robust viral infection in Tc1 polarizing conditions appear sufficient for full-fledged T cell differentiation.

We speculate that once dividing, the newly generated daughter effector CD8 T cells and possibly also newly dividing daughter CD4 T cells can use their high affinity LFA-1 to engage ICAMs on neighboring T cells rather than on DCs. These ICAMs likely provide the daughter effector T cells generated by initially primed OT-I T cells, as well as their descendent T cells with the ability to confine their locally secreted IL-2 and IFNγ to nearby daughter T cells as was demonstrated in several studies (*16, 64, 65*). In this scenario, effector daughter T cells with a shared TCR, if remaining nearby each other, will likely generate homotypic clusters with recently divided sister T cells using their high affinity LFA-1 and both ICAM-1 and ICAM-2 (*66*). A clue for the physiological importance of this scenario is provided by studies on IL-2 signaling between effector T cells with shared TCRs which augments their quorum sensing, (*67, 68*). In parallel, non-cognate DCs may provide additional costimulatory TCR independent signals, such as CD40L-CD40 especially to rapidly dividing effector T cells (*62*). It is also possible that CD8 T_cm_ generated from naïve T cells during vaccination of both control LNs and LNs deficient in DC ICAM-1 acquire additional ICAM-1-facilitated pro-survival signals from non APCs in the form of growth factors, in particular IL-7 and IL-15 (*69–71*). Consistent with this possibility, our high-resolution single cell transcriptomic analysis of memory CD8 generated in vaccinated LNs deficient in DC ICAM-1 revealed remarkably conserved memory signatures of these lymphocytes compared to memory CD8 T cells generated in similarly vaccinated LNs of control mice.

In conclusion, the assumption that DC ICAM-1 provides essential co-stimulatory signaling via T cell LFA-1 during CD8 T cell proliferation in strong Th1 polarizing conditions must be revisited. Our present work raises the intriguing possibility that LFA-1-ICAM-1 strengthened CD4 and CD8 T-DC synapses may be critical only for specialized cases of immunizations, most probably when the Ag presenting DCs are rare and remote from each other. In that scenario the likelihood of cooperative T cell triggering by multiple adjacent pMHC-presenting DCs is diminished and therefore any reduced stability of individual highly stable ICAM-1 dependent DC-T contacts cannot be compensated by highly frequent weaker DC-T contacts. Such a scenario may prevail when a small number of DCs are loaded with high levels of antigens. Nevertheless, in most viral infections, which involve either direct drainage via lymphatics or migratory DCs carrying viral antigens into draining LNs (*72, 73*), the DCs either die or cross-present the viral antigen to numerous resident DCs, each presenting a low number of pMHC complexes. Our current results predict that during such conditions, frequent Ag triggered synapses, if properly timed, can deliver differentiation responses as potent as rare strong ICAM-1 stabilized DC-T contacts. Future studies on additional infection models and other types of T cell differentiation processes should reveal when ICAM-1-mediated strengthening of T-DC synapses is mandatory for optimal T cell differentiation into effector and memory T cells. Such insights should help in the future design of genetically modified DCs for improved vaccination against viral infections.

## MATERIALS AND METHODS

### Study design

The aim of this study was to establish the role of ICAM-1 specifically expressed on DCs in early CD8 activation and differentiation into effector and memory T cells. For this, we generated the first described conditional DC ICAM-1 KO mice, and transferred WT transgenic CD8 T cells with defined specificity to the model antigen ovalbumin (OVA). T cell activation, proliferation and differentiation following vaccination and viral infection was assessed using a combination of flow cytometry and single cell 10x RNAseq analysis. The study was complemented by DC-T conjugate quantitative flow cytometric analysis, supplemented by immunofluorescence and imaging flow cytometry. In addition, functional T cell readouts such as *in vivo* cytotoxicity, migration, and responses in various recall immune responses were applied. The number of mice per experimental group and the number of repetitions of the experiments are indicated in individual figure legends.

### Mice

To generate the ICAM-1^fl/fl^ mice, single-stranded oligonucleotides (LoxP sites as well as the restriction site for screening purposes) were co-injected with optimized single guide RNAs (sgRNAs) and Cas9 mRNA into mice embryos derived from the hybrid CB6F1, where the embryos are F2 from male C57BL/6 x female BALB/c. These LoxP sites were inserted using CRISPR-Cas9 upstream of the first exon and downstream of the second exon of ICAM-1 (Fig. S1A). This location was chosen in order to avoid known regulatory regions (*74, 75*) and ensure no residual protein, as the internal exons are all in frame (*76*). Mice generated by this procedure (Founder, F0) were genotyped via PCR for upstream and downstream insertion. Double positive mice were validated via sequencing with primers to regions outside of the LoxP sites (Fig. S1B). The F1 generation of the founder (F0) double positive mice were similarly genotyped for the 2 LoxP sites. The double positive F1 mice had both insertions on the same allele, and this was also validated via sequencing (not shown). For backcrossing to the C57BL/6 background, we used the speed congenic approach (Fig. S1C) (DartMouse, the mouse speed congenic facility at The Geisel School of Medicine at Dartmouth). This technique was performed by analyzing PCR amplified regions of simple sequence length polymorphism (SSLP) markers between the recipient and donor strains: offspring with the highest number of markers showing the recipient genome across all chromosomes were chosen for the next generation(*77*). After completing a fourth round of speed congenics, the top male was identified as being 97% C57BL/6 (Fig. S1C).

The generation of ICAM-1^fl/fl^ mice is described in the supplementary information. DC ICAM-1 KO mice were generated by crossing ICAM-1^fl/fl^ CD45.2 male mice with female CD45.2 mice carrying the cell type specific Cre-recombinase under the CD11c-Cre transgene (*35*). Pups were validated via PCR for LoxP site insertions, ICAM-1 and Cre+ or Cre-expression. OT-I (C57BL/6-Tg(TcraTcrb)1100Mjb/J) TCR transgenic mice harboring OVA-specific CD8+ T cells were purchased from the Jackson Laboratory. OT-I tdTomato mice were kindly provided by Z. Shulman (Weizmann Institute, Israel). All mice used were 8-12 weeks old, and both males and females were used for all experiments. All mice procedures were performed in agreement with the institutional guidelines for animal care and all experimental protocols were approved by the Institutional Animal Care and Use Committee (IACUC) of the Weizmann Institute.

### Immunizations and cell isolations

OT-I T cells were isolated from spleens of CD45.1 mice and intraveneously injected (1×10^7^ cells) either unlabeled or following CFSE labeling (5 µM, Molecular Probes) into recipient CD45.2 DC ICAM-1 KO mice and their control counterparts. 18 hrs later the recipient mice received a subcutaneous intrafootpad injection of either αDEC-205:OVA (0.05 µg/footpad, kindly provided by R. Dahan, Weizmann Institute) plus αCD40 antibody (10 µg/footpad, clone FGK4.5/FGK45, BioXCell) or MVA-OVA (1×10^7^ IU, (*53*)).

For DC isolation, LNs were sliced into small pieces and digested for 30 min at 37°C in collagenase D (1 mg/ml, Roche). For DC-T conjugate analysis by flow cytometry and ImageStream, popliteal LNs were digested under mild conditions in Isocove’s modified Dulbecco’s medium (Sigma-Aldrich) supplemented with Liberase-TL (100μg/ml, Roche) and DNase I (100μg/ml, Roche), and incubated with frequent agitation at 3°C for 20 min. For recall response, mice were challenged by subcutaneous intrafootpad injection of 40 µg OVA (Sigma-Aldrich) emulsified in 2.5 mg/ml CFA (Difco).

### Flow cytometry

Cells were stained with the following Abs: allophycocyanin (APC)–conjugated anti-CD19 (clone 6D5; catalog no. 115512), B220 (clone RA3-6B2; catalog no. 103212), CD62L (clone MEL-14; catalog no. 104412), CD25 (clone PC61; catalog no. 102011), I-A/I-E (clone M5/114.15.2; catalog no. 107614), CD54 (ICAM-1, clone YN1/1.7.4; catalog no. 116120), alexa fluor 647-conjugated anti-CD102 (ICAM-2, clone 3C4; catalog no. 105612), phycoerythrin (PE)–conjugated anti-CD54 (YN1/1.7.4; catalog no. 116108), CD3 (clone 145-2C11; catalog no. 100307), CD40 (clone 3/23; catalog no. 124610), TNFα (clone MP6-XT22; catalog no. 12-7321-81, eBioscience), CD69 (clone H1.2F3; catalog no. 104507), CD8α (clone 53-6.7; catalog no. 100707), CD11a (LFA-1, clone M17/4; catalog no. 12-0111-83, eBioscience), fluorescein (FITC)-conjugate anti-CD8α (clone 53-6.7; catalog no. 100706), Siglec H (clone 551; catalog no. 129603), pacific blue-conjugated anti-CD45.1 (clone A20; catalog no. 110722), granzyme B (clone GB11; catalog no. 515408), alexa fluor 488-conjugated anti-CD80 (clone 16-10A1; catalog no. 104715), brilliant violet 421-conjugated anti-CD11c (clone N418; catalog no. 117343), PE/Cy7-conjugated anti-CD11b (clone M1/70; catalog no. 101216), CD169 (clone 3D6.112; catalog no. 142411), IFNγ (clone XMG1.2; catalog no. 505826), CD11c (clone N418; catalog no. 117318), APC/Cy7-conjugated anti-CD19 (clone 6D5; catalog no. 115529), CD44 (clone IM7; catalog no. 103028), CD3 (clone 17A2; catalog no. 100221), I-A/I-E (clone M5/114.15.2; catalog no. 107628), PerCP/Cy5.5-conjugated anti-Ly6C (clone HK14; catalog no. 128012), CD86 (clone GL-1; catalog no. 105028), CD11b (clone M1/70; catalog no. 101228), CD8α (clone 53-6.7; catalog no. 100734), CD3 (clone 17A2; catalog no. 100218), brilliant violet 510-conjugated anti-I-A/I-E (clone M5/114.15.2; catalog no. 107635), brilliant violet 605-conjugated anti-CD103 (clone 2E7; catalog no. 121433). All antibodies were purchased from Biolegend, unless otherwise indicated. Intracellular IFNγ, granzyme B and TNFα staining was performed using the Cytofix/Cytoperm Kit (BD Biosciences) according to the manufacturer’s instructions. Before intracellular staining, cells were restimulated for 1.5 hrs with PMA (50 ng/ml) and ionomycin (500 ng/ml) (Sigma-Aldrich) and then treated with Brefeldin A solution (Biolegend, 1:1,000) and Monensin (Biolegend, 1:1000), for 1.5 hours. Flow cytometry was performed on CytoFlex flow cytometer (Beckman Coulter) and analyzed using FlowJo software v.10.7.1 (Tree Star). cDC1 were gated as CD19^-^ CD11c^high^MHCII^+^CD8^+^CD11b^-^CD103^+^, cDC2 as CD11c^high^MHCII^+^CD8^-^CD11b^+^CD103^-^, moDCs as CD11c^+^MHCII^+^Ly6C^+^CD11b^+^CD103^-^, pDCs as CD11c^int^MHCII^int^B220^+^SiglecH^+^CD19^-^, B cells as CD11c^-^MHCII^+^CD19^+^ and SCS macrophages as CD11c^int^MHCII^+^CD169^+^CD11b^+^CD19^-^. For T_cm_ sorting, popliteal LN single cell suspensions were stained and gated as CD3^+^CD8^+^CD45.1^+^CD62L^+^CD44^+^. Sorting was performed on an Aria III sorter (BD Biosciences). Conjugates were imaged using imaging flow cytometer (ImageStreamX mark II: Amnis Corp, Seattle, WA) and analyzed using the manufacturer’s image analysis software (IDEAS 6.2; Amnis Corp).

### Immunofluorescence staining

Popliteal LNs fixed for 2 hrs in 1% paraformaldehyde followed by 30% sucrose exchange for 2 days were embedded in OCT (Tissue-Tek). Sections of 10μm thickness were generated with a cryostat (Leica) and dehydrated in acetone prior to freezing. Before staining, sections were rehydrated in PBS and blocked with PBS with 0.05% Tween-20 and 3% BSA for 1 hr. Then, slides were stained with Alexa Fluor 647-conjugated anti CD11c (clone N418; catalog no. 117314, Biolegend) in 1% BSA in PBS/T overnight at 4°C. Slides were washed and stained shortly with Hoechst33342 (1:20,000, Thermo Fisher Scientific). Sections were mounted with mounting medium (Sigma-Aldrich) and confocal imaging was performed using a Zeiss LSM 880 confocal microscope. The acquired images were processed using ImageJ software. DC-T clusters were defined as T cells in close proximity with DCs (i.e. less than 1 µm). Quantification of the clusters was performed by analysis of up to 6 different sections from differently treated LNs.

### *In vivo* killing assay

Recipient mice (CD45.2) were transferred with CD45.2 OT-I splenocytes and immunized with αDEC-205:OVA plus αCD40 as described in previous sections. 72 hrs later, splenocytes were isolated from non-immunized CD45.1 WT mice. Half of these target cells were pulsed with OVA_257-264_ peptide (SIINFEKL, 1 µg per 10^6^ cells) for 2 h at 37°C. Unpulsed and pulsed cells were incubated, respectively, with either low (0.5 µM) or high (5 µM) doses of CFSE. 1×10^7^ of each group of target cells were co-transferred by intravenous injection into the immunized mice. The differential clearance of both target cell populations by endogenously generated OT-I CTLs was evaluated by flow cytometry one day later. Specific killing was defined as (1-%CFSE^high^/%CFSE^low^) x100.

### Droplet-based scRNA-seq

CD45.1 OT-I LNs cells isolated from two DC ICAM-1 KO or control mice 72 hrs post immunization with αDEC-205:OVA plus αCD40, as described in previous sections, were independently stained with TotalSeq™-B0303 anti-mouse Hashtag 3 Antibody (Biolegend, cat: 155835) according to the manufacturer’s instructions and with fluorophore-conjugated antibodies for cell sorting (anti-CD45.1-PB, anti-CD8-PerCP/Cy5.5 and anti-CD3-APC). Next, the samples were sorted using Sony HS800S sorter and pooled right before running the 10x Genomics single-cell 3’ v. 3.1 assay. Single cell RNA-seq libraries were prepared using the Chromium Single Cell Controller (10x Genomics, Pleasanton, CA). Briefly, 30,000 cells were loaded into the 10X genomics channel. Single cells were partitioned into droplets with gel beads in the Chromium Controller. After emulsions were formed, barcoded reverse transcription of RNA took place. This was followed by cDNA amplification, fragmentation and adapter and sample index attachment, all according to the manufacturer’s recommendations. Libraries were pooled together and sequenced on one flow cell of an Illumina NovaSeq, with paired end reads as follows: read 1, 28 nt; read 2, 90 nt; index 1, 10 nt; index 2, 10 nt.

### Computational Analysis

#### Pre-processing of droplet (10X) scRNA-seq data

Sequencing data from the 10x Genomics libraries was demultiplexed, aligned to the mm10 genome and UMI-collapsed using the count function of the Cellranger toolkit (version 6.0.1) provided by 10X Genomics. Further demultiplexing of the hashtag oligos was performed using the Seurat R package (version 4.0.3) (*78*) according to the instructions in the vignette: https://satijalab.org/seurat/articles/hashing_vignette.html. Doublets (cells with two antibody tags) and negatives (no antibody tag) were removed and the singlets assigned to the relevant samples. Cells with more than 13% mitochondrial UMI counts, cells with less than 300 UMI counts and cells in the top or bottom 1% of genes or UMI counts were removed from the analysis (function FilterCells). After the filtration the number of cells was 1648, 1616, 2173 and 1740 for the control and DC^DICAM-1^ duplicates respectively.

### scRNA-seq data analysis

All further single cell analyses were performed using the Seurat R package (version 4.0.3) using default parameters unless otherwise stated. The data was normalized using the function SCTransform (*79*). This function replaces the previous commands NormalizeData, ScaleData, and FindVariableFeatures. Dimensionality reduction was performed using PCA (RunPCA). The functions RunUMAP and FindNeighbors were applied using 12 PCs. The clustering was performed with a resolution of 0.6. Ten clusters were detected, two of them were removed from the analysis, one because it had a high number of cells expressing mitochondrial genes and the other one because of low *Cd8a* expression. Differential gene expression was calculated using the functions FindAllMarkers or FindMarkers with the default statistical test Wilcoxon rank-sum test. Multiple hypothesis testing correction was performed by controlling the false discovery rate (FDR)(*80*) using the R function p.adjust.

### Differential expression of signature gene-sets

T effector and T memory cells signature genes sets were obtained from (*81*). The first 20 genes with the lowest FDR and with detected gene expression were selected for the signatures. For the T effector signature gene set we used: *Gzmb, Tmsb4x, Lgals1, Crip1, Atp5j, Rrm2, Actr2, Ly6a, S100a10, Stmn1, Top2a, Hmgb2, Plac8, Atp5k, Actb, Calm3, S100a6, Pycard, H2afz, Txn1* and for T memory signature: *Wnt4, Klf6, Npy1r, Tcf7, Cd22, Ecm2, Bach2, Slfn5, Gmip, Il7r, Ccdc78, Kcnmb4, H2-Q4, Mc1r, Zscan2, Btg1, Slx1b, H2-DMb2, Btg2, Disc1*. The co-inhibitory cell signature gene set was obtained from (*48*). The human gene symbols were converted into their mouse orthologues using the MGI Batch Query in the Mouse Genome Informatics website (http://www.informatics.jax.org/batch). The genes used for this signature were *Lag3, Lilrb4a, Pdcd1, Tigit, Klra3, Lair1, Klrc3, Ptger4, Havcr2, Btla, Cd200, Klrc1, Cd160, Klrd1, Klra7, Cd244a and Cd274*. The mean of the normalized, log transformed and scaled gene expression value of all the signature genes was calculated per cell and used for the violin plots and statistical analysis. Violin plots were drawn using the ggplot2 R package. Signatures were compared between OT-I cells isolated from DC^DICAM-1^ and control (for the entire samples and for each cluster separately) using a linear mixed-effects model, with genotype as a fixed factor and sample ID as a random factor. p-values were corrected for multiple comparisons by FDR. Effect size between genotypes was calculated by Cohen’s D. All mixed models were run in R (v. 4.1.0), using the package ‘lme4’ and ‘lmerTest’.

### Testing for changes in cell type proportions

The number of cells in each cluster was compared between genotypes using a binomial mixed-effects model (logistic regression), with genotype as a fixed factor and sample ID as a random factor. p-values were corrected for multiple comparisons by FDR. Mixed models were run in R (v. 4.1.0), using the package ‘lme4’ and ‘lmerTest’. A positive log-odds ratio indicates a higher proportion of cells in DC^DICAM-1^ samples.

### Statistical analysis

Analysis was performed using Prism 9.0c version (\**P* < 0.05, \*\**P* < 0.01, \*\*\**P* < 0.001, \*\*\*\**P* < 0.0001 and n.s. represents nonsignificant). All n numbers represent independent biological repeats and points in graphs indicate individual mice. In bar and dot graphs, bars indicate means and error bars indicate SD. Significance was assessed by Student’s two-tailed unpaired t test to determine the significance of the difference between means of two groups. One-way ANOVA was used to compare means among three or more independent groups. Tukey’s post hoc tests were used to compare all pairs of treatment groups when the overall *P* value was < 0.05. Littermates and sex-matched animals were used whenever possible.

## Supplementary Materials

### Supplementary Figures

Fig S1. Generation of CD11c-Cre:ICAM-1^fl/fl^ mice.

Fig S2. Comparable ICAM-2 expression on cDCs of control and DC-specific ICAM-1 KO mice after vaccination with αDEC-205:OVA plus αCD40.

Fig S3. Titration of in vivo OT-I priming and differentiation by increasing doses of αDEC-205:OVA and a fixed dose of αCD40 Ab antibody.

Fig. S4. T cell homing and activation to popliteal LNs in DC-specific ICAM-1 KO

Fig. S5. Comparable expression of co-stimulatory molecules on cDCs of control and DC-specific ICAM-1 KO mice.

Fig. S6. Identification and gating of DC-T cell conjugates in mice immunized with αDEC-205:OVA plus αCD40.

Fig. S7. Comparable OT-I T cell proliferation following αDEC-205:OVA plus αCD40 vaccination in DC ICAM-1 depleted and control mice.

Fig. S8. Intracellular IFNγ, granzyme B and TNFα staining of OT-I T cells following αDEC-205:OVA plus αCD40 immunization.

Fig. S9. Single cell analysis of OT-I T cells isolated from popliteal LNs of immunized DC ICAM-1 depleted and control mice.

Fig. S10. Comparable early activation of transferred OT-I T cells in MVA-OVA infected LNs of DC ICAM-1 depleted and control mice.

Fig. S11. OT-I T cell proliferation and intracellular IFNγ staining of the OT-I T cells following MVA-OVA infection of DC ICAM-1 depleted and control mice.

### Supplementary Tables

Table S1. Differential expression analysis between different subsets of OT-I T cells isolated from immunized DC ICAM-1 depleted and control mice.

Table S2. Differential expression analysis between OT-I T cells isolated from immunized DC ICAM-1 depleted and control mice.

## Acknowledgments

We thank Profs. Steffen Jung and Ziv Shulman and the members of the Alon group for their critical review of the manuscript. We also thank Prof. Wolfgang Kastenumuller (University of Würzburg) and Dr. Rony Dahan for providing reagents. We thank Drs. Rebecca Haffner-Krausz and Shifra Ben-Dor for invaluable help in the design and generation of the new transgenic mice used in this study, and Drs. Dena Leshkowitz, Gil Stelzer, and Ron Rotkopf for help in the bioinformatic and statistical analysis. We also thank Dr. Hadas Keren-Shaul from the Sandbox Unit of the WIS Life Sciences Core Facilities for help with the scRNA-Seq experiments and Dr. Ziv Porat from the Life Sciences Core Facilities for imaging flow cytometry data analysis.

## Funding

Israel Science Foundation (791/17, RA) (1587, MB))

Minerva Stiftung (RA, MB)

Moross Integrated Cancer Center (RA, MB)

Helen and Martin Kimmel Institute for Stem Cell Research (RA, MB)

German Israeli Foundation (I-1470-412.13/2018) (RA)

Israel Cancer Research Fund (19-109-PG) (RA)

EU Horizon 2020 Research and Innovation Program (Ri-boMed 857119) (RA)

Yeda-Sela Center for Basic Research (RA)

Meyer Henri Cancer Endowment (RA)

William and Marika Glied and Carol A. Milett (RA)

## Author contributions

Conceptualization: RA, AS

Methodology: RA, AS

Investigation: AS, SWF, SK, ND, EP-K, AG

Single cell computational analysis: NW, EF, MB

Supervision: RA, MB

Writing – original draft: RA, AS, MB

Writing – review & editing: RA, AS, SWF, MB

## Data and materials availability

All data are available in the main text, in the supplementary materials, or will be deposited in a public database. R scripts enabling the main steps of the bioinformatics analysis will be made available upon request.

