## Supplementary Figures for "Dendritic cell ICAM-1 strengthens immune synapses but is dispensable for effector and memory responses"

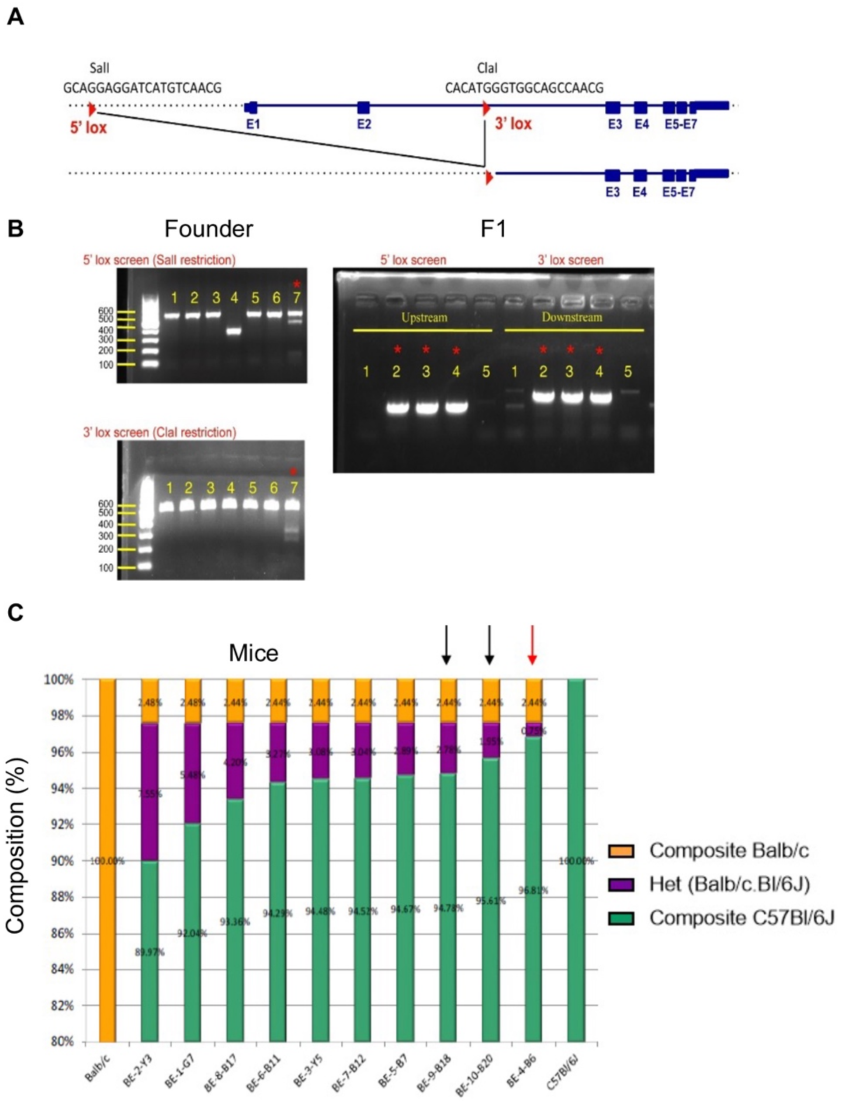


**Figure S1. Generation of CD11c-Cre:ICAM-1$^{fl/fl}$ mice**

**(A)** Design of conditional ICAM-1 locus. Lox sites (red triangles) were inserted using CRISPR-Cas9 upstream of the first exon and downstream of the second exon of ICAM-1. Guide sequences are shown above each lox site, as well as the restriction site introduced with the Lox for screening purposes. **(B)** PCR of founder (F0) and F1 mice. 16 F1 mice were generated, with one founder male positive for both insertions (#7, left panels). Eleven F1 mice were screened, and eight were found positive for both insertions, indicating that both were on the same allele. Three such representative mice are shown (right panel) **(C)** Graphs documenting the percentages of the heterozygosity and homozygosity in representative animals (DartMouse, Dartmouth Geisel School of Medicine). The SNP is color-coded, green: C57BL/6J, purple: heterozygous (C57BL/6J x Balb/c), orange: Balb/c. The data shows that the animals with the least heterozygosity among those submitted with this panel of samples are 4-B6 (red arrow) followed by 9-B18 and 10-B20 (black arrows). These are the animals that were recommended and taken for next generation breeding.

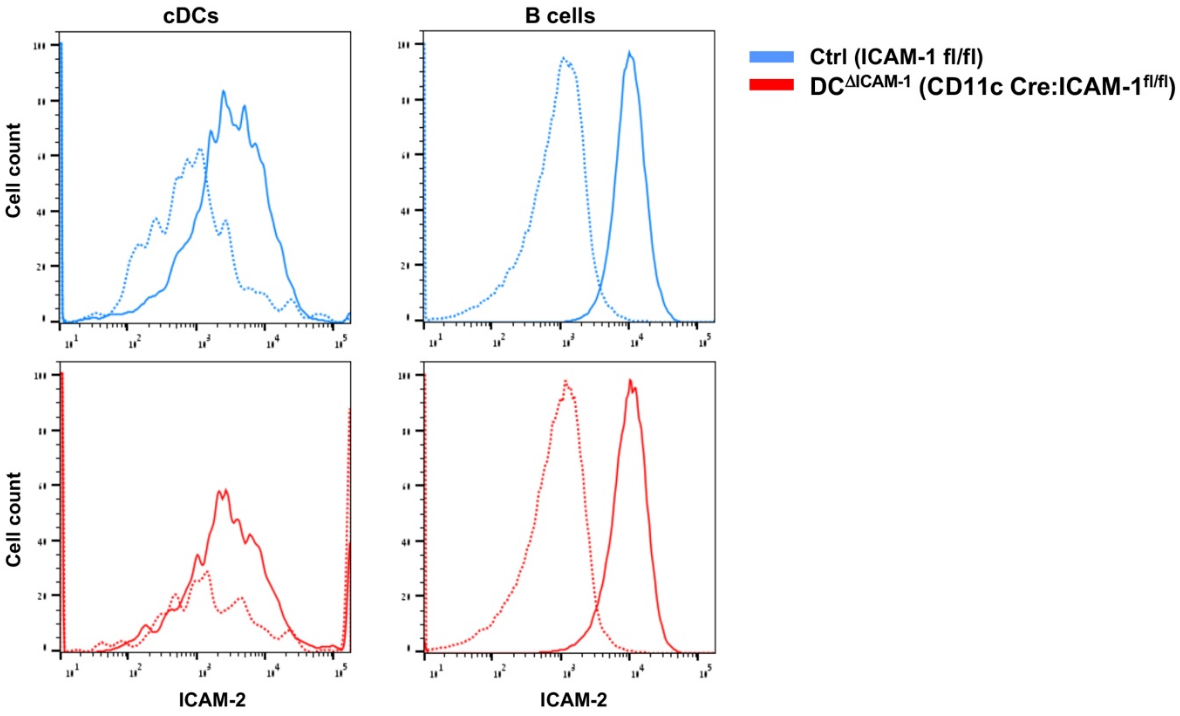


**Figure S2. Comparable ICAM-2 expression on cDCs of control and DC-specific ICAM-1 KO mice after vaccination with αDEC-205:OVA plus αCD40**

Control (blue, solid line) or $DC^{\Delta ICAM\text{-}1}$ (red, solid line) mice were injected intrafootpad with αDEC-205:OVA (0.05 μg/footpad) plus αCD40 (10 μg/footpad) and 24 hrs later cDCs and B cells were recovered from popliteal LNs and analyzed for ICAM-2 expression. Isotype mAb controls are shown in dashed lines. (*n* = 3/group).

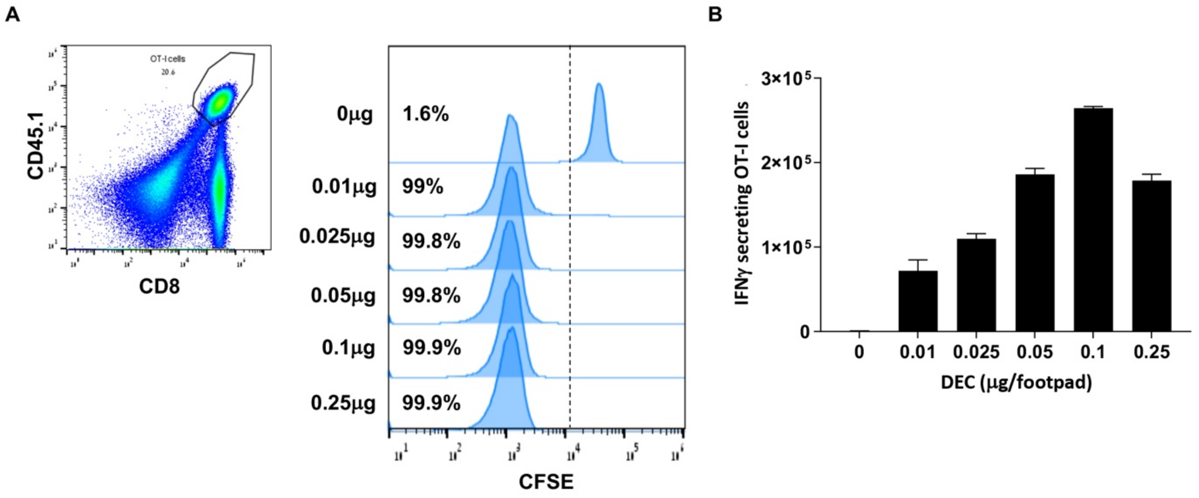


**Figure S3. Titration of *in vivo* OT-I priming and differentiation by increasing doses of αDEC-205:OVA and a fixed dose of αCD40 antibody**

**(A)** Mice were transferred with CFSE-labeled OT-I T cells, 18 hrs later underwent intrafootpad immunized with indicated doses of αDEC-205:OVA plus αCD40, and 72 hrs later CFSE profiles (and percentage CFSE-diluted cells) were determined. Dot plot represents gating of transferred OT-I T cells. **(B)** Bar graphs represent intracellular IFNγ staining of OT-I cells 72 hrs after αDEC-205:OVA plus αCD40 immunization. (*n* = 3/group)

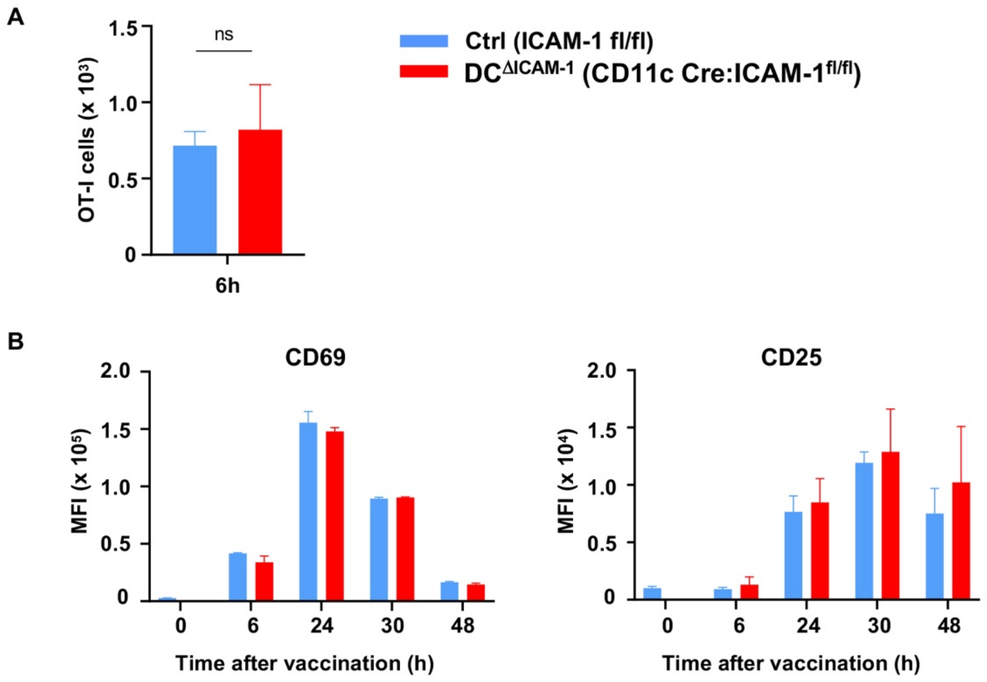


**Figure S4. T cell homing and activation to popliteal LNs in DC-specific ICAM-1 KO mice is identical to control mice**

**(A)** Control (blue) or $DC^{\Delta ICAM-1}$ (red) mice were injected with αDEC-205:OVA plus αCD40 and 6 hrs later homing of transferred OT-I T cells to the popliteal LNs was determined. (*n* = 3/group, unpaired *t* test). **(B)** MFI of CD69 and CD25 expression by the transferred OT-I cells in control (blue) and $DC^{\Delta ICAM-1}$ (red) mice at indicated time-points following αDEC-205:OVA plus αCD40 immunization. (*n* = 3/group)

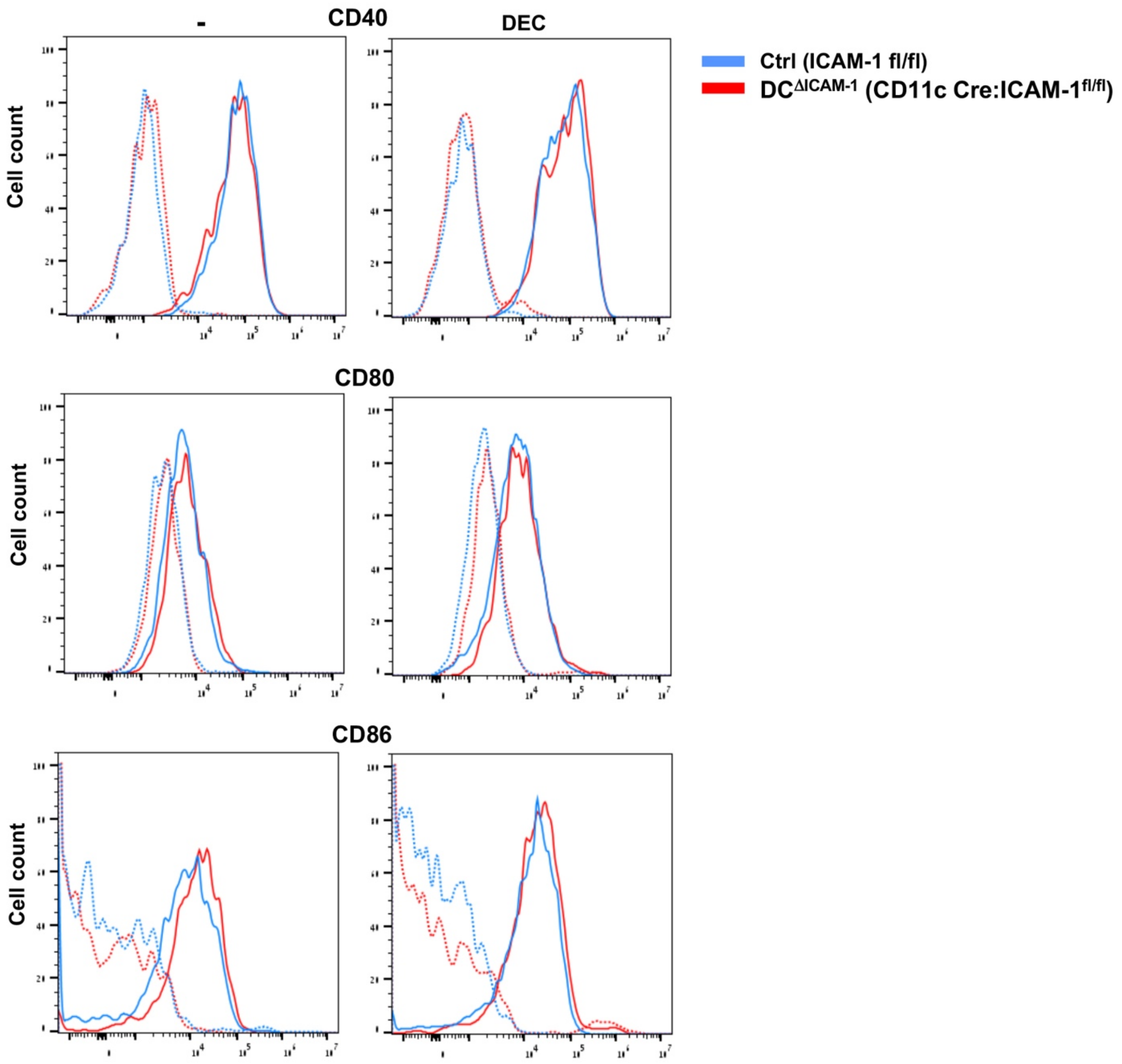


**Figure S5. Comparable expression of co-stimulatory molecules on cDCs of control and DC-specific ICAM-1 KO mice**

Control (blue, solid line) or $DC^{\Delta ICAM\text{-}1}$ (red, solid line) mice were injected intrafootpad with αDEC-205:OVA (0.05 μg/footpad) plus αCD40 (10 μg/footpad) (DEC) or PBS (-) and 24 hrs later cDCs were recovered from popliteal LNs and analyzed for CD40, CD80 and CD86 expression. Isotype mAb controls, are shown in dashed lines. (*n* = 3/group)

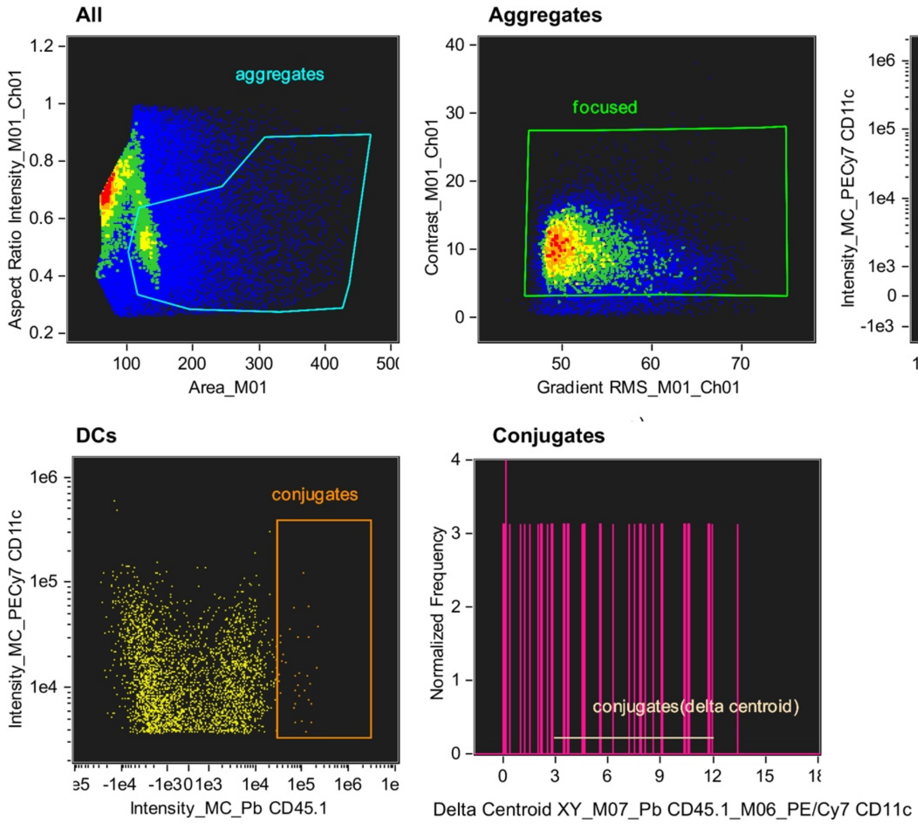


**Figure S6. Identification and gating of DC-T cell conjugates in mice immunized with αDEC-205:OVA plus αCD40**

Mice were immunized with αDEC-205:OVA plus αCD40 18 hrs after transfer of CD45.1 OT-I cells, and DC–T cell conjugates were imaged using imaging flow cytometry 24 hrs after vaccination. Aggregates of cells (excluding single cells) were gated using the area (the number of microns squared) and aspect ratio (the minor axis divided by the major axis) features on the bright-field channel. Focused cells were selected using the gradient RMS and contrast features (which measure the sharpness quality of an image) on the bright-field image. Aggregates that contain DCs were selected based on the intensity of staining for CD11c and MHCII. Out of DC containing cell aggregates, cell conjugates containing OT-I T cells were further selected based on the intensity of staining for CD45.1. DC-T cell conjugates were further analyzed using Delta Centroid XY feature that calculated the distance between the centers of stainings of CD11c (M 6-mask of channel 6) and CD45.1 (M7-mask of channel 7). Conjugates with intermediate delta centroid XY values demonstrating a synapse connection between DC and OT-I T cell were selected.

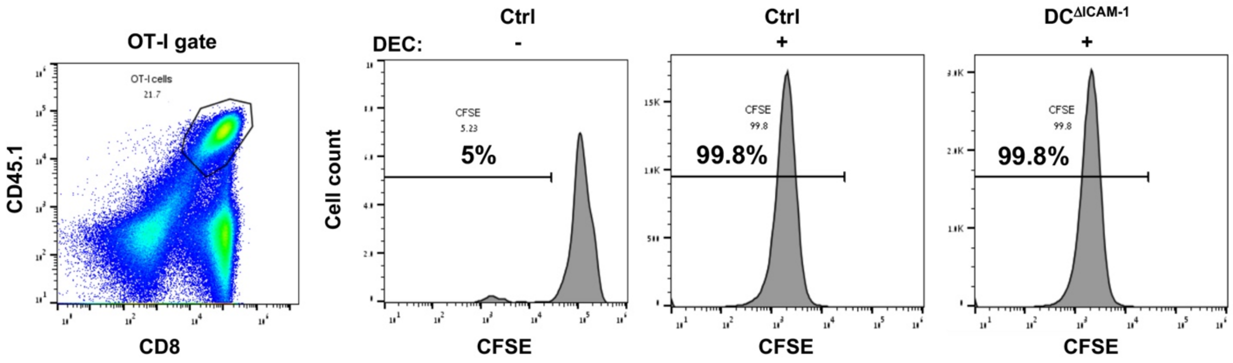


**Figure S7. Comparable OT-I T cell proliferation following αDEC-205:OVA plus αCD40 vaccination in DC ICAM-1 depleted and control mice**

CFSE-labeled OT-I CD45.1 T cells were transferred into control and DC$^{\Delta ICAM-1}$ mice and 18 hrs later the mice were immunized with αDEC-205:OVA plus αCD40 (+) or PBS (-) . 72 hrs following the immunization, proliferation of OT-I T cells was analyzed by CFSE dilution. Dot plots represent gating for OT-I transferred cells; histograms represent CFSE dilution and numbers indicate the percentage of CFSE-diluted cells.

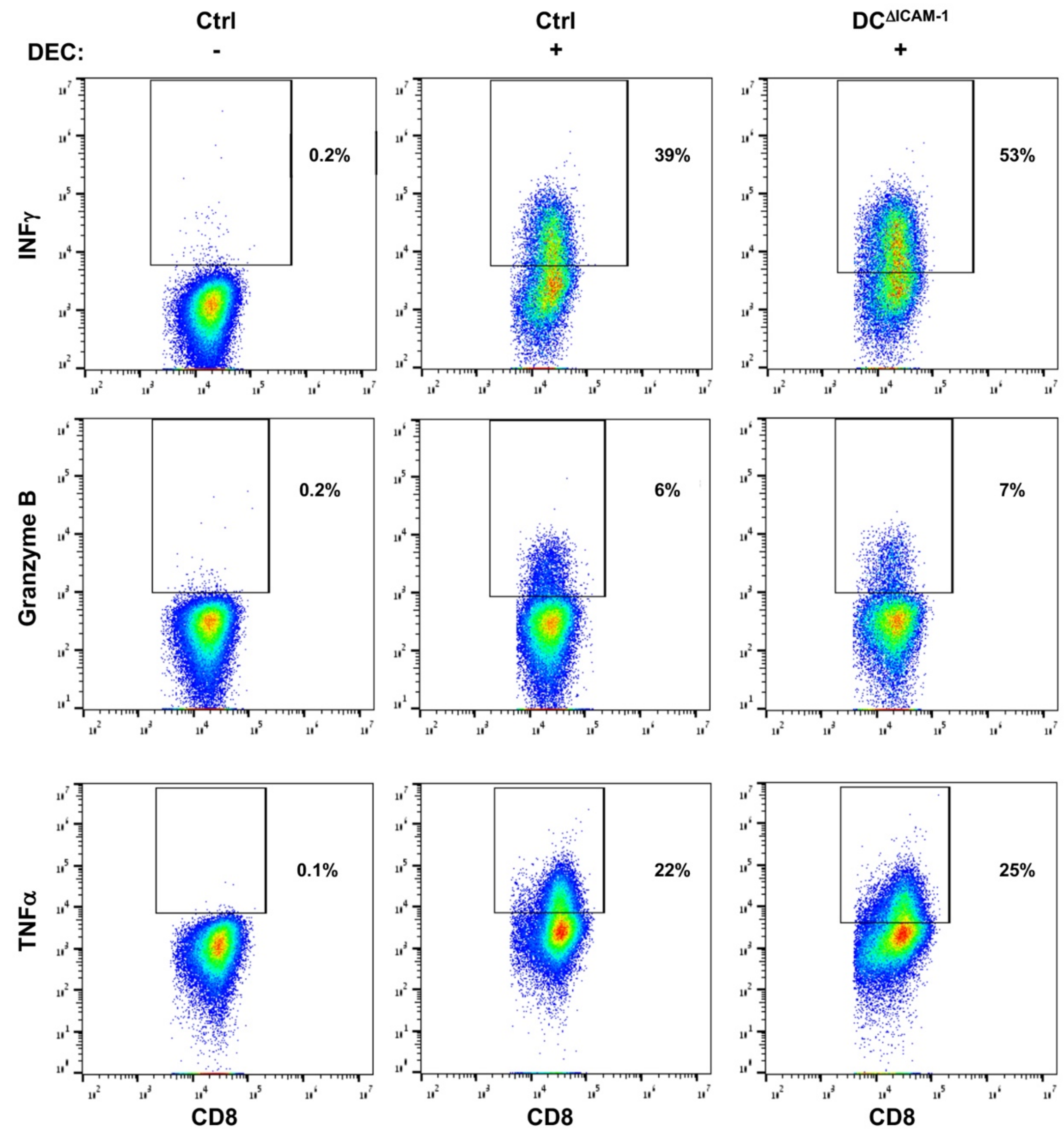


**Figure S8. Intracellular IFNγ, granzyme B and TNFα staining of OT-I T cells following αDEC-205:OVA plus αCD40 immunization**

Dot plots represent gating of IFNγ, granzyme B and TNFα positive OT-I cells isolated from popliteal LNs 72 hrs following intrafootpad vaccination with αDEC-205:OVA plus αCD40 (+) or PBS (-) of control and $DC^{\Delta ICAM\text{-}1}$ mice.

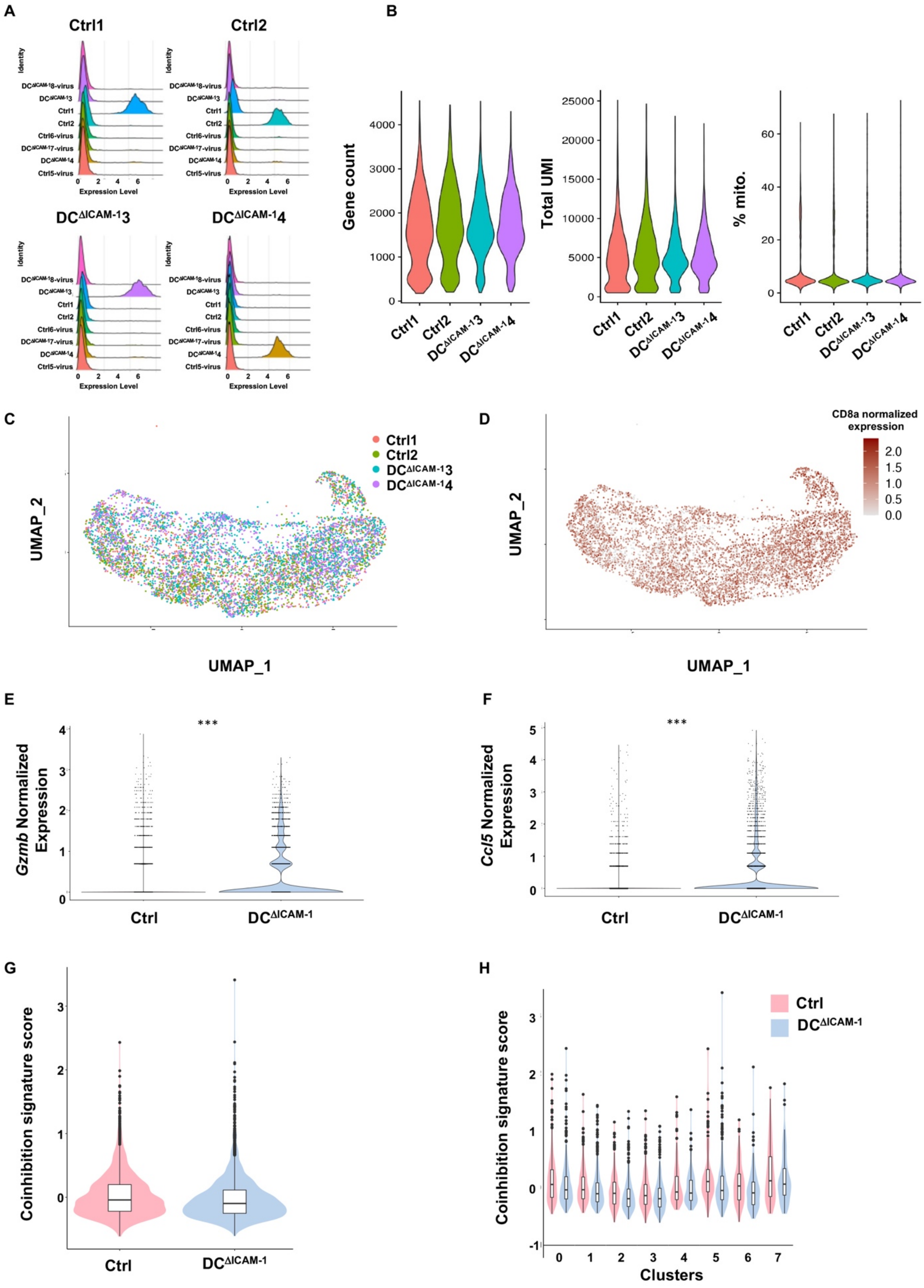

A
Ctrl1
Ctrl2
DC$^{ΔICAM-1}$3
DC$^{ΔICAM-1}$4
Identity
DC$^{ΔICAM-1}$8-virus
DC$^{ΔICAM-1}$3
Ctrl1
Ctrl2
Ctrl6-virus
DC$^{ΔICAM-1}$7-virus
DC$^{ΔICAM-1}$4
Ctrl5-virus
Expression Level
B
Gene count
Total UMI
% mito.
Ctrl1
Ctrl2
DC$^{ΔICAM-1}$3
DC$^{ΔICAM-1}$4
C
Ctrl1
Ctrl2
DC$^{ΔICAM-1}$3
DC$^{ΔICAM-1}$4
UMAP_2
UMAP_1
D
CD8a normalized expression
UMAP_2
UMAP_1
E
***
*Gzmb* Normalized Expression
Ctrl
DC$^{ΔICAM-1}$
F
***
*Ccl5* Normalized Expression
Ctrl
DC$^{ΔICAM-1}$
G
Coinhibition signature score
Ctrl
DC$^{ΔICAM-1}$
H
Ctrl
DC$^{ΔICAM-1}$
Coinhibition signature score
Clusters

**Figure S9. Single cell analysis of OT-I T cells isolated from popliteal LNs of immunized DC ICAM-1 depleted and control mice**

**(A)** Count distribution of the most highly expressed hashtag oligonucleotide (HTO) *(82)* per group to identify the sample origin. Four different samples of OT-I T cells isolated from popliteal LNs 72 hrs following intrafootpad vaccination with αDEC-205:OVA plus αCD40 of DC-specific ICAM-1 KO mice ($DC^{\Delta ICAM-1}$) (*n*=2) or control mice (*n*=2) were identified and used for subsequent analysis. **(B)** Quality control metrics of OT-I T cells isolated from popliteal LNs of DC-specific ICAM-1 KO mice (*n*=2) or control mice (*n*=2) immunized as in (A) are shown. Number of genes (left), number of UMIs (middle) and percentage of mitochondrial genes (right) for each sample are shown. **(C,D)** Uniform Manifold Approximation and Projection (UMAP) for Dimension Reduction of 6693 single cells (points). (**C**) Demultiplexed cell subsets, colored by HTO, *n*=2 samples per genotype. **(D)** colored by normalized expression of the *CD8a*. **(E,F)** Expression of statistically significant upregulated effector T cell genes in OT-I T cells from $DC^{\Delta ICAM-1}$ compared to control mice. Violin plot showing the distribution of *Gzmb* expression **(E)**, or *Ccl5* **(F**) isolated from control or DC-ICAM-1 depleted or control mice (*n*=2 mice per group).****p* value < 0.0001, *Wilcoxon test*. (**G,H**) Expression of co-inhibitory T cell signatures in OT-I T cells. Violin plots showing the distribution of mean expression values of co-inhibitory signature (17 genes) *(51)* expressed in control or DC-ICAM-1 depleted mice (*n*=2 samples per group). **G**, all OT-I cells, **H**, OT-I T cell subsets. No significant statistical changes were found (FDR). Bar showing median and box interquartile range.

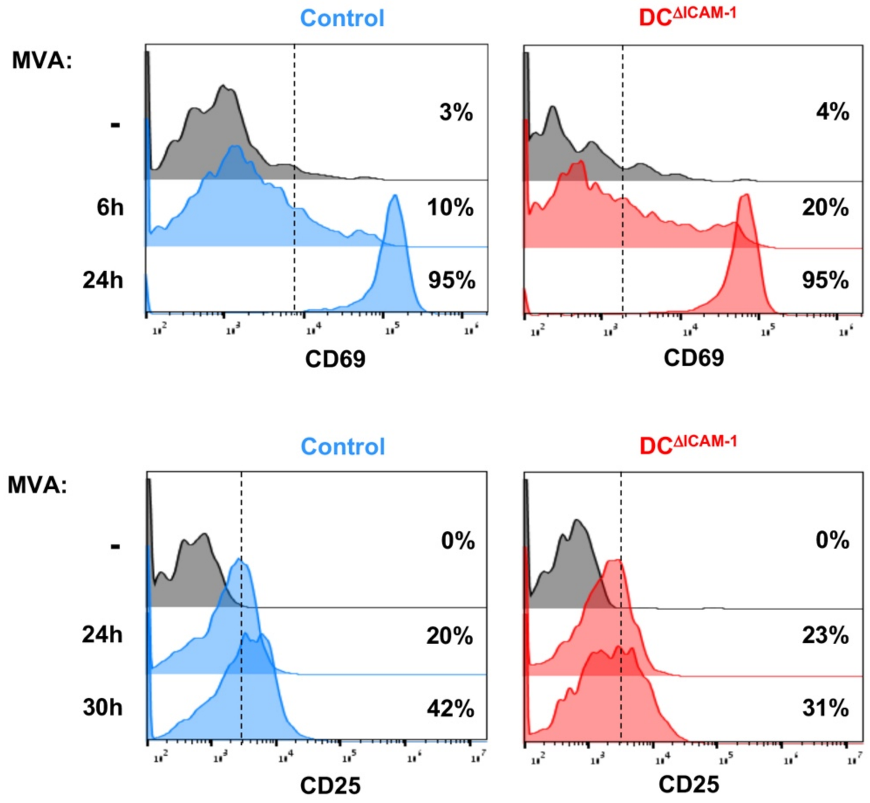


**Figure S10. Comparable early activation of transferred OT-I T cells in MVA-OVA infected LNs of DC ICAM-1 depleted and control mice**

Induction of CD69 (top) or CD25 (bottom) on intravenously transferred OT-I cells in popliteal LNs of either control (blue) and $DC^{\Delta ICAM\text{-}1}$ (red) mice 6 and 24 or 24 and 30 hrs (for CD69 and CD25, respectively) following intrafootpad infection of MVA-OVA (times indicated) or PBS (-). (*n* = 3/group)

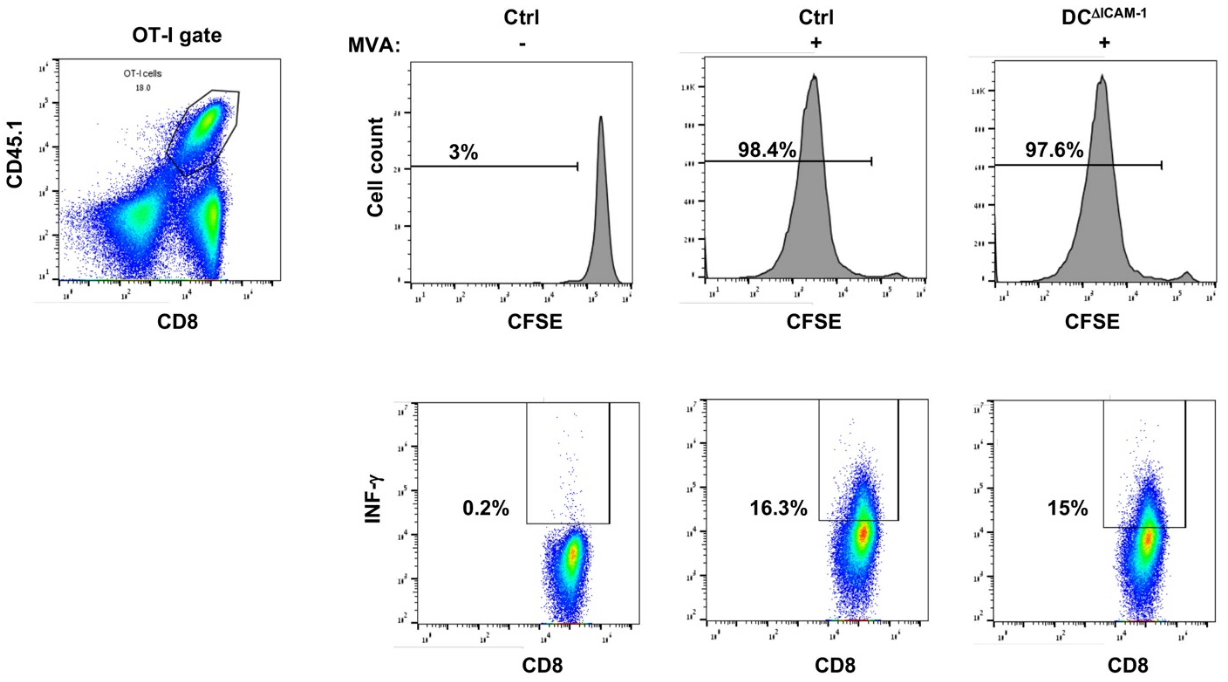


**Figure S11. OT-I T cell proliferation and intracellular IFNγ staining of the OT-I T cells following MVA-OVA infection of DC ICAM-1 depleted and control mice**

CFSE-labeled OT-I CD45.1 T cells were transferred into control and DC$^{\Delta ICAM-1}$ mice, and 18 hrs later the mice were immunized with MVA-OVA (+) or PBS (-). 72 hrs following the infection, proliferation of OT-I T cells was analyzed by CFSE dilution. Dot plots represent gating for OT-I transferred cells; histograms represent CFSE dilution and numbers indicate the percentage of CFSE-diluted cells. Dot plots in the lower panel represent gating of IFNγ positive OT-I cells isolated from popliteal LNs 72 hrs following intrafootpad infection with MVA-OVA.

### Supplementary Tables

**Table S1. Differential expression analysis between different subsets of OT-I T cells isolated from immunized DC ICAM-1 depleted and control mice.** Related to Fig. 5 and Fig. S9. Differentially expressed genes of the different clusters of OT-I T cells (cluster 0-7, see tabs). The genes are ranked by adjusted *p* values. Statistically significant genes are colored in black and the non-significant are colored in grey.

**Table S2. Differential expression analysis between OT-I T cells isolated from immunized DC ICAM-1 depleted and control mice.** Related to Fig. 5 and Fig. S9. Differentially expressed genes per cluster (cluster 0-7, see tabs), or bulk analysis of the data (Tab "bulk"). The genes are ranked by adjusted *p* values. Statistically significant genes are colored in black and the non-significant are colored in grey.
